# Wnt5A Signaling Regulates Gut Bacterial Survival and T cell Homeostasis

**DOI:** 10.1101/2022.07.18.500401

**Authors:** Soham Sengupta, Suborno Jati, Shreyasi Maity, Malini Sen

## Abstract

In light of the demonstrated antagonism of Wnt5A signaling toward the growth of several bacterial pathogens, it was important to study the influence of Wnt5A on gut resident bacteria, and its outcome. Here we demonstrate that in contrast to inhibiting the survival of the established gut pathogen *Salmonella enterica*, Wnt5A clearly promotes the survival of the common gut commensals *Enterococcus faecalis* and *Lactobacillus rhamnosus* within macrophages through a self-perpetuating Wnt5A-Actin axis. A Wnt5A – Actin axis furthermore regulates the subsistence of the natural bacterial population of the Peyer’s patches, as is evident from the diminution in the countable bacterial colony forming units therein through the application of Wnt5A signaling and actin assembly inhibitors. Wnt5A dependency of the gut resident bacterial population is also manifested in the notable difference between the bacterial diversities associated with the feces and Peyer’s patches of Wnt5A heterozygous mice, which lack a functional copy of the Wnt5A gene, and the wild type counterparts. Alterations in gut commensal bacterial population resulting from either the lack of a copy of the Wnt5A gene or inhibitor mediated attenuation of Wnt5A signaling correlate with significantly different ratios of regulatory vs. activated CD4 T cells associated with the Peyer’s patches. Taken together, our study reveals the importance of Wnt5A signaling in shaping the gut commensal bacterial population and the T cell repertoire linked to it, thus unveiling a crucial control device for the maintenance of gut bacterial diversity and T cell homeostasis.

**Significance Statement:** Gut commensal bacterial diversity and T cell homeostasis are crucial entities of the host innate immune network. Yet molecular details of host directed signaling pathways that sustain the steady state of gut bacterial colonization and T cell activation remain unclear. Here we describe the protective role of a Wnt5A-Actin axis in the survival of several gut bacterial commensals, and its importance in shaping gut bacterial colonization and the associated T cell repertoire. This study opens up new avenues of investigation into the role of the Wnt5A-Actin axis in protection of the gut from dysbiosis related inflammatory disorders.

## Introduction

Wnt5A belongs to the19 – member family of glycoprotein ligands that initiate signaling upon binding to the Frizzled and ROR/Ryk families of transmembrane cell surface receptors. While Frizzleds, about 12 in number, resemble the heterotrimeric G protein coupled receptors, ROR1, ROR2 and Ryk resemble tyrosine kinases (1–3). Wnt - Frizzled/ROR/Ryk signaling has long been known to be crucial for tissue patterning and organism development (4–7). Through the course of research across different disciplines it has now become clear that Wnt signaling also features as a prominent player in the regulation of infection, inflammation and immunity in different modes that are to a large extent context dependent (8–11).

Wnt signaling operates as two major pathways – β -catenin dependent (canonical) and β -catenin independent (non-canonical). Wnt directed canonical signaling pathways almost invariably involve the transcriptional activation of β-catenin – Lef/Tcf responsive genes, whereas Wnt directed non-canonical signaling pathways often operate independent of β-catenin. Although Wnt5A signaling represents the non-canonical mode of Wnt signaling as per classification, due to the existing homology among the different Wnt cognate receptors and the presence of common Wnt signaling intermediates, overlap between Wnt5A signaling with the canonical mode of Wnt signaling is not infrequent (12–14).

In compliance with the involvement of Wnt5A signaling in organelle trafficking and cell polarity (7, 15, 16) we demonstrated along with others the role of Wnt5A in actin remodeling and phagocytosis of foreign matter including microbes (8, 17, 18). We also demonstrated that Wnt5A regulated actin organization directs the killing of microbial pathogens through the generation of autophagosome like moieties within macrophages (19). These findings led us to examine the interrelation between Wnt5A signaling and the commensal bacteria that form part and parcel of human physiology.

Among other anatomical niches within the mammalian host, the Peyer’s patches of the gut are specially noted for harboring phagocytes and a multitude of commensal bacteria that are crucial for the maintenance of gut immune homeostasis (20–23). Quite naturally, loss or change in the bacterial population leading to alterations in bacterial diversity is linked with the development of several autoimmune disorders (24). Although an interrelation between Wnt expression and intestinal commensal bacterial colonization has been suggested (25), much remains unknown about whether or how Wnt signaling in the steady state controls the intestinal bacterial population thereby influencing gut immunity.

In the current study, using mouse macrophages, isolated mouse Peyer’s patches, and a mouse model, we examined if Wnt5A signaling is involved in the maintenance of gut microbial diversity and T cell activation profile. Our experimental results reveal that Wnt5A signaling aids in the survival of a multitude of commensal bacteria, which could be crucial for the preservation of gut CD4 T cell homeostasis.

## Results

### Wnt5A signaling promotes uptake and survival of common gut commensal bacteria by macrophages

In order to evaluate the influence of Wnt5A signaling on the cellular uptake and survival of gut commensal bacteria, we initially focused on two common gut commensals, *Enterococcus faecalis* and *Lactobacillus rhamnosus*, using the macrophage as a representative phagocyte of the gut mononuclear phagocyte system (26). The *Enterococcus faecalis* strain used for experiments described here was initially identified through screening of the mouse cecum resident bacteria by Wnt5A mediated augmented internalization into macrophages and later validated by 16S sequencing (**Fig. S1A and B**). Wnt5A mediated augmented uptake of *E. faecalis* by macrophages (both RAW264.7 and mouse peritoneal macrophages) was subsequently confirmed by enumerating the colony forming units (cfu) from independent uptake experiments using recombinant Wnt5A (**Fig. 1A**). *Lactobacillus rhamnosus*, purchased commercially and furthermore validated by 16S sequencing (**Fig. S1C**) was also internalized in higher numbers by the same macrophages in response to added Wnt5A, when compared to the corresponding vehicle (PBS) control (**Fig. 1B**). Augmented uptake of both *E. faecalis* and *L. rhamnosus* in the Wnt5A activated macrophages was also demonstrated separately by microscopy (**Fig. 1C**and **D**). Continued incubation of the Wnt5A activated infected macrophages for several hours following bacterial uptake (0 time point) indicated that increased uptake correlates with increased survival/proliferation of the internalized bacteria with the course of time (**Fig. 1E**). That Wnt5A signaling facilitates the survival of these commensals was furthermore corroborated by the observed attenuation in bacterial cfu through application of inhibitors of the Wnt5A signaling intermediates Disheveled (Dsh) (27, 28) and Rac1 (29, 30), and anti-Frizzled5 antiserum (31) to the Wnt5A activated bacteria harboring macrophages, which were left to incubate for 3 (T3) or 6 (T6) hr following infection (**Fig. 2A** and **B)**. Inhibitor mediated decrease in bacterial counts in the absence of added Wnt5A (PBS samples) clearly suggested that Wnt5A signaling at the steady state is involved in bacterial survival. A requirement of steady state Wnt5A signaling for the survival of *E. faecalis* and *L. rhamonosus* was validated by the loss of bacterial cfu in the bacteria infected RAW264.7 macrophages where Wnt5A level was reduced by Wnt5A siRNA transfection (**Fig. 2C**). Reduction of Wnt5A was evident from immunoblotting and subsequent densitometry (**Fig. 2D**). Wnt3A reduction did not have any significant effect on the survival of *E. faecalis* and *L. rhamnosus* in macrophages as demonstrated by similar experiments indicating that there is considerable specificity in the interrelation between these gut commensals and Wnt5A signaling (**Fig. 2E** and **F**). The pro-survival effect of Wnt5A on *E. faecalis* and *L. rhamnosus* was in stark contrast to its antagonism toward the growth of the gut pathogen *Salmonella enterica* in both RAW264.7 and peritoneal macrophages (**Fig. 2G**). The non-pathogenic nature of *E. faecalis* and *L. rhamnosus*, both of which formed biofilms, was supported by their vancomycin sensitivity. *S. enterica*, the gut pathogen used as a reference demonstrated vancomycin resistance (**Fig. S2**).

**Fig. 1.**
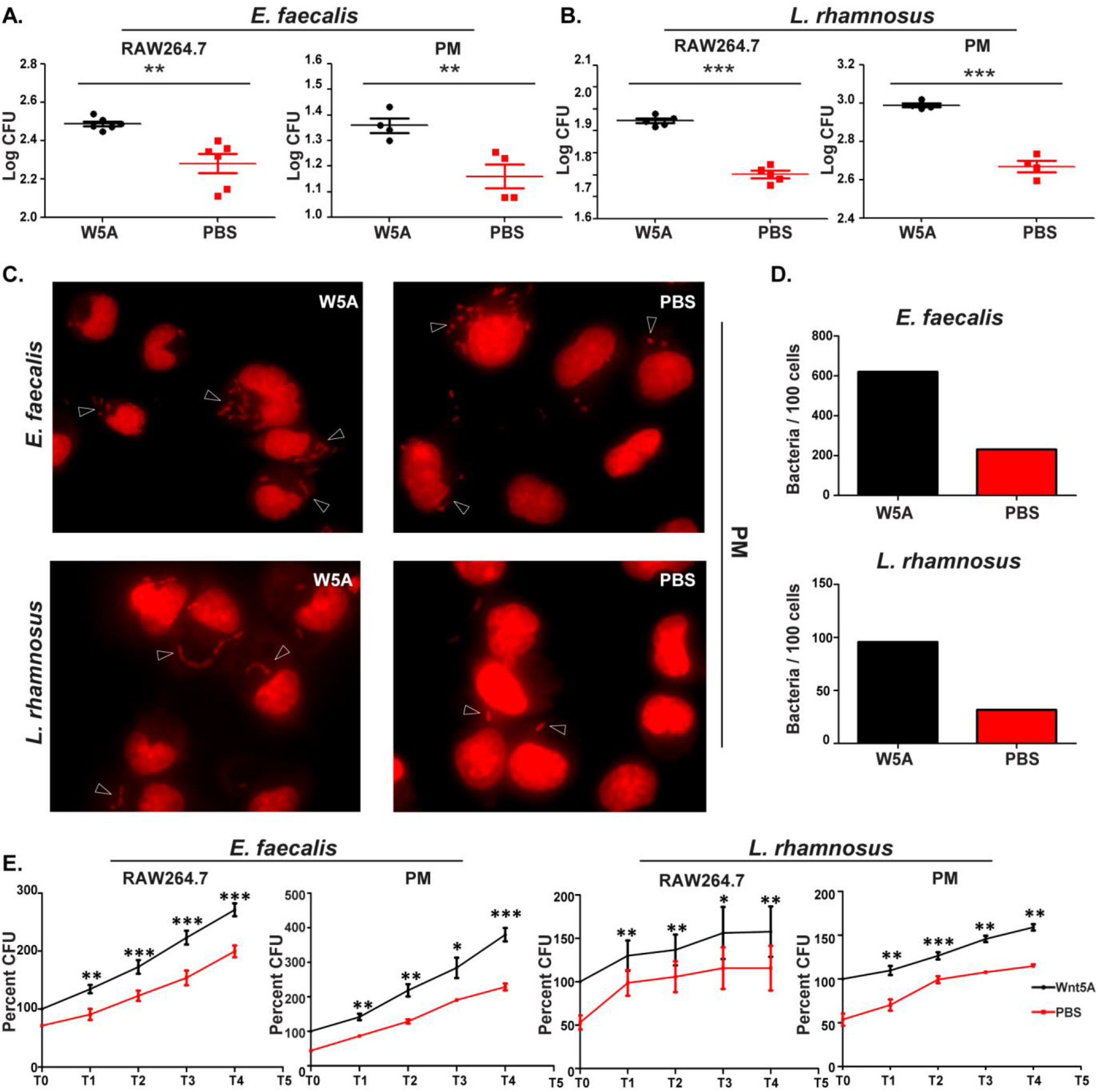
Wnt5A aided internalization and survival of bacterial commensals. (A) Pretreatment of RAW 264.7 (n=6) and peritoneal macrophages (PM) (n=4) with rWnt5A for 6hours led to higher internalization of *E. faecalis* after 2 hr infection at MOI 10, as evident from cfu. (B) Similar increase in internalization of *L. rhamnosus* (enumerated by cfu) following infection (MOI 20) of rWnt5A pretreated RAW264.7 (n=5) and peritoneal macrophages (n=4). (C) and (D) *E. faecalis* (C) and *L. rhamnosus* (D) internalization in peritoneal macrophages (rWnt5A vs. PBS pretreated) as observed by fluorescence microscopy of Propidium Iodide stained cells. (E) Enumeration of number of bacteria/100 cells as observed in different microscopy fields (F) Wnt5A mediated survival/proliferation of internalized *E. faecalis* and *L. rhamnosus* (n=3-4) as enumerated by cfu, 1 – 4 hr post infection. “n” represents number of experiments unless stated otherwise. Data is represented as mean ± SEM. Significance was represented by * in the following manner: * p≤ 0.05, ** p≤ 0.005, *** p≤ 0.0005.

**Fig. 2.**
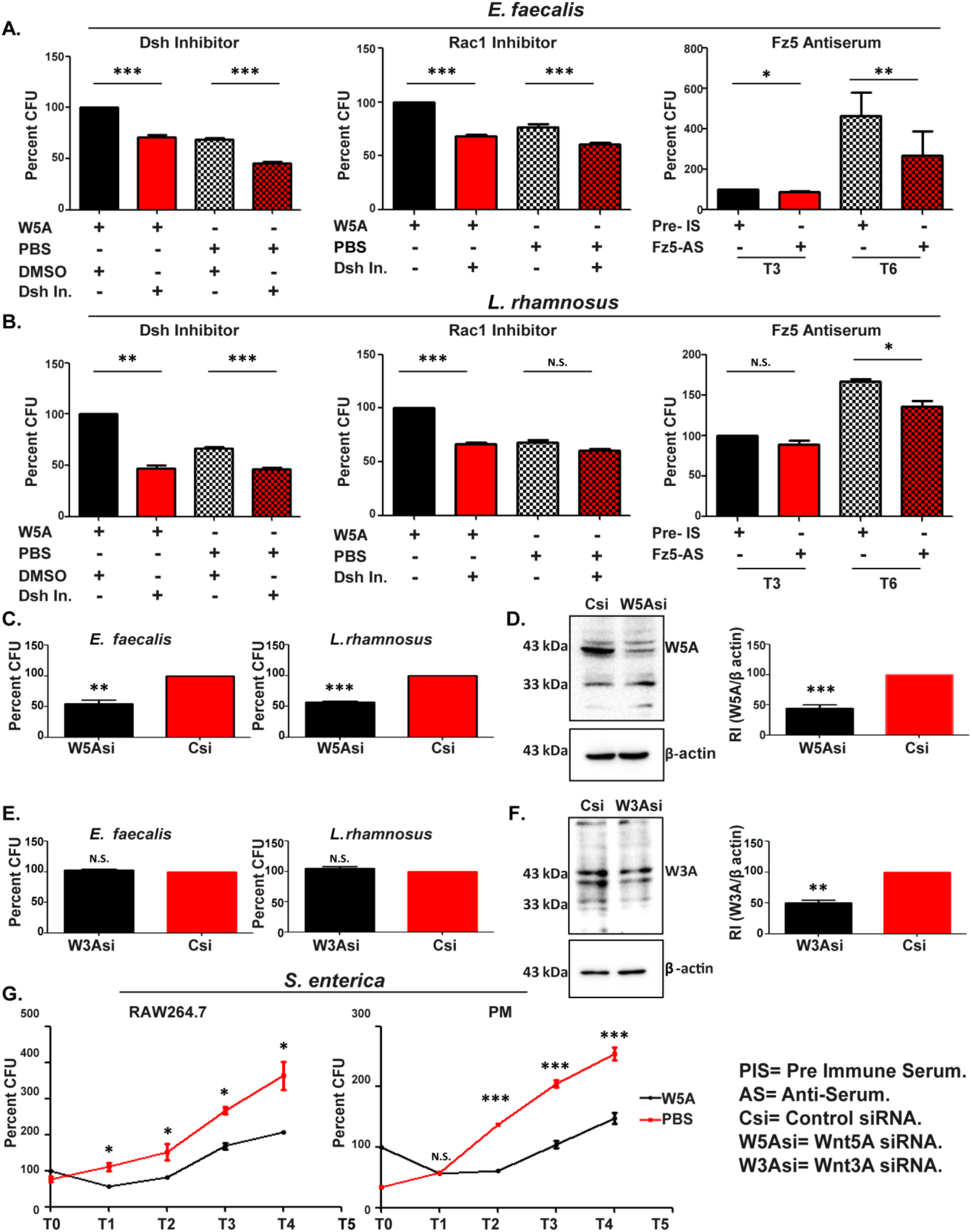
Inhibition of Wnt5A signaling blocks survival of commensal bacteria. (A) and (B): Application of inhibitors of Wnt5A signaling intermediates Dsh and Rac-1 for 3 hr, as well as blocking of Fz5 receptor by application of antiserum for 3 and 6 hr post infection by *E. faecalis* (MOI 10 for 1 hr) (A) and *L. rhamnosus* (MOI 20) (B) led to cfu reduction in infected RAW264.7 cells (n=3-6). (C) Knockdown of Wnt5A by siRNA post internalization of *E. faecalis* and *L. rhamnosus* led to decrease in bacterial survival as enumerated by cfu (n=4). (D) Immunoblot and densitometry showing decrease of Wnt5A in Wnt5A siRNA transfected cells as compared to control. (E) Downregulation of Wnt3A by siRNA did not have significant effect on bacterial survival (n=4). (F) Immunoblot and densitometry confirming siRNA mediated Wnt3A depletion. (G) Upregulation of Wnt5A signaling opposed the survival of internalized *S. enterica* (n=3 for both RAW264.7 and Peritoneal macrophages). Data represented as mean ± SEM and p≤ 0.05 was considered significant statistically. Significance was represented by * in the following manner: * p≤ 0.05, ** p≤ 0.005, *** p≤ 0.0005. “N.S.” denotes non-significant.

### A Wnt5A-Actin axis supports the survival of commensal bacteria

In light of the demonstrated association of actin dynamics with Wnt5A signaling (9, 17, 32), we investigated if Wnt5A assisted survival of the commensal bacteria is linked with actin assembly. Indeed, Wnt5A depletion induced reduction in *E. faecalis* and *L. rhamnosus* survival in macrophages was clearly associated with diminution in actin assembly as demonstrated by confocal microscopy and subsequent ImageJ (intensity) analysis of phalloidin stained cells (**Fig. 3A and B**). The necessity of actin assembly for both *E. faecalis* and *L. rhamnosus* survival, and the involvement of Wnt5A therein, was separately demonstrated by actin assembly inhibitor mediated loss of bacterial cfu that was considerably recovered by the addition of rWnt5A to the inhibitor treated *E. faecalis* and *L. rhamnosus* harboring macrophage cultures during a 3-hour incubation period following infection. The significant gain in bacterial cfu by the addition of rWnt5A in the absence of actin assembly inhibitors furthermore confirmed the pro-survival effect of the Wnt5A-Actin axis (**Fig. 3C**). Interestingly, *E. faecalis* or *L. rhamnosus* internalization (T0: 0 hr) and continued incubation of the infected macrophages (T3, T4: 3 hr, 4 hr) resulted in increased secretion of Wnt5A (**Fig. 3D**) that correlated with near perfect maintenance of macrophage actin assembly (**Fig. 3E**). The intactness of actin assembly in the *E. faecalis* or *L. rhamnosus* infected macrophages was clearly distinct from the almost dilapidated state of actin in the *S. enterica* infected macrophages. These observations point toward a coordinated self-sustaining circuit of Wnt5A signaling (autocrine/paracrine), actin assembly and bacterial survival in the *E. faecalis* or *L. rhamnosus* infected, but not *S. enterica* infected macrophages (**Fig. S3**). Such a concept is in agreement with the observed blockade in *E. faecalis* or *L. rhamnosus* survival by the administration of anti-Frizzled5 antiserum and other WNt5A signaling inhibitors to the bacteria harboring macrophages (Fig. 2).

**Fig. 3.**
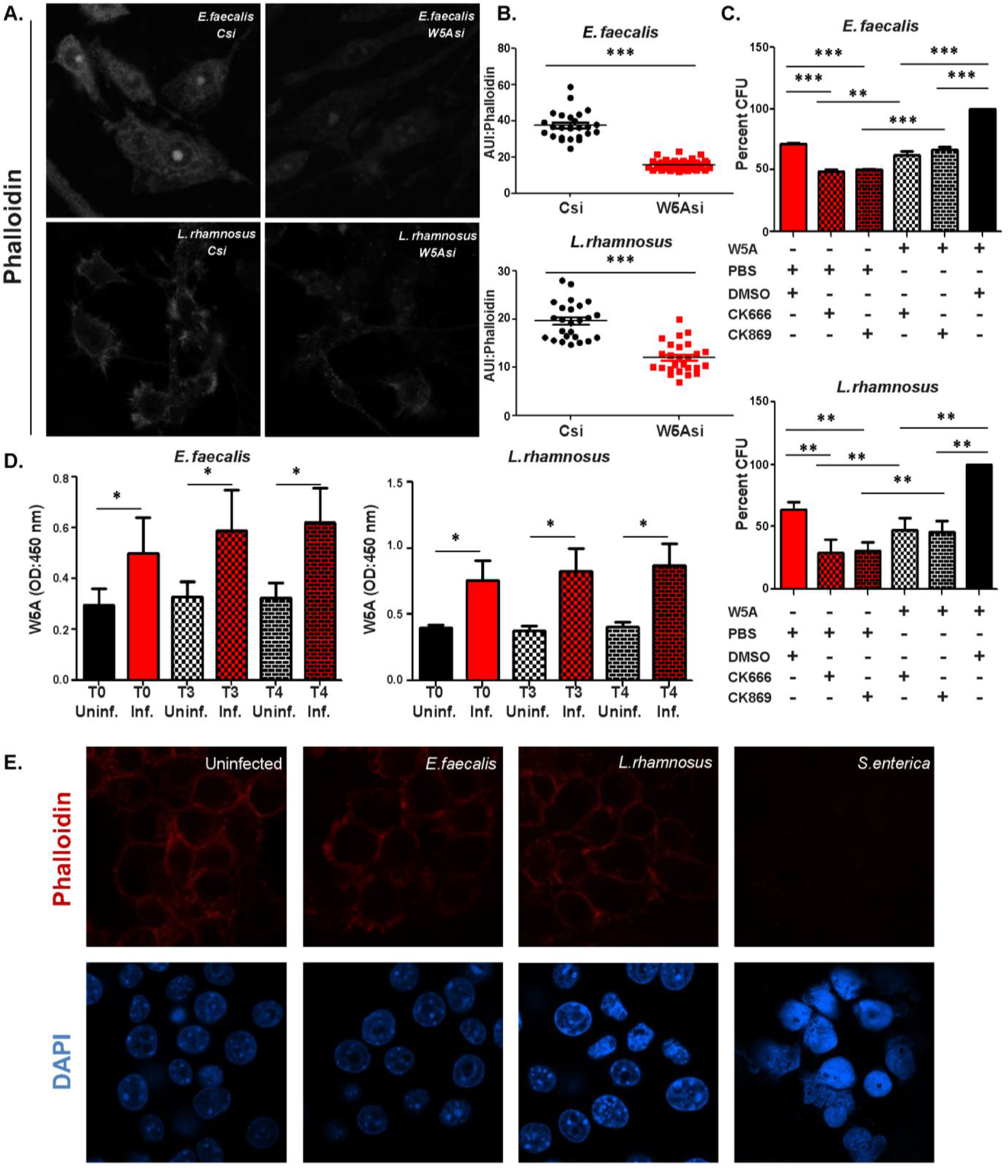
Wnt5A-Actin axis supports survival of commensal bacteria. (A) Wnt5A downregulation by siRNA led to decreased actin accumulation in RAW264.7 cells infected with *E. faecalis* and *L. rhamnosus* as noted from confocal microscopic images of Phalloidin stained cells. Images represented in monochrome. (B) Enumeration of Arbitrary Unit of Intensity (AUI) of phalloidin staining as measure of actin assembly (n=25 cells). (C) Application of actin polymerization inhibitors CK666 and CK 869 inhibited survival of internalized *E. faecalis* and *L. rhamnosus* that was considerably recovered by addition of rWnt5A. rWnt5A without actin assembly inhibitors promoted maximum survival (n=4). (D) Infection of RAW264.7 cells by *E. faecalis* and *L. rhamnosus* increased secretion of Wnt5A 3 and 4 hr (T3 and T4) post infection (T0) as observed by indirect ELISA (n=5). (E) Infection with *E. faecalis* and *L. rhamnosus* maintained actin assembly as seen from Phalloidin staining of RAW264.7 cells. The pathogen *S. enterica* severely hampered actin organization. Data represented as mean ± SEM and p≤ 0.05 was considered as significant statistically. Significance was represented by * in the following manner: * p≤ 0.05, ** p≤ 0.005, *** p≤ 0.0005.

### Wnt5A signaling regulates the growth/colonization of commensal bacteria in the mouse gut

In view of the fact that the Peyer’s patches of the small intestine constitute a major hub for gut resident commensal bacteria and phagocytes (21, 22, 33–35), we investigated the influence of Wnt5A signaling on the growth/colonization of commensal bacteria within the Peyer’s patches. Having demonstrated that the Peyer’s patches isolated from BALB/c mouse gut express and secrete Wnt5A by both immunoblotting and ELISA (**Fig. 4A** and **B**), we also confirmed the existence of intracellular bacteria therein (depicted by arrowheads), by microscopy of cultured adherent cells of the Peyer’s patches after propidium iodide (PI) staining (**Fig. 4C**). Additionally, the presence of *Enterococcus sp*. and *Lactobacillus sp*. in the Peyer’s patches was ascertained by both qPCR of RNA isolated from harvested tissue and PCR of genomic DNA isolated from cultured adherent cells (**Fig. 4D** and **Fig. S4**). Subsequently, we demonstrated that modulation of Wnt5A signaling by the administration of IWP-2 and Dsh inhibitor affects bacterial survival in the Peyer’s patches. **Fig. 4E** depicts significant reduction in the BHI/MRS agar grown bacterial colonies harvested from the total cells of the Peyer’s patches following incubation of the cells for 6 hr separately with IWP-2 and Dsh inhibitor. DMSO was used as vehicle control in these assays. **Fig. S5** depicts inhibition of Wnt5A signaling by the Dsh inhibitor and blockade of Wnt5A secretion by IWP-2. Inhibitor mediated alteration in the number of countable bacterial colonies was also reflected in the adherent cells of the Peyer’s patches (**Fig. 4F**), which constitute both CD11c+/CD11b+/ CD11c+CD11b+ and CD11c- CD11b- phagocytes (26). The presence of PI stained intracellular bacteria (red: depicted by arrowhead) already demonstrated in **Fig. 4C** was additionally verified in the context of CD11b+ (green) phagocytes by confocal microscopy (**Fig. 4G**). The same inhibitor IWP-2 also blocked the survival of *E. faecalis* and *L. rhamnosus* after their internalization (T0) in the adherent cells of the Peyer’s patches during a 3 hr (T3) incubation period (**Fig. 4H**). These adherent cells were pretreated with antibiotic for removal of endogenous bacteria. In accordance with the inherent association of actin assembly with Wnt5A signaling, we furthermore found that application of actin assembly inhibitors led to significant reduction in the number of BHI/MRS agar grown bacterial colonies harvested from the isolated cells of the Peyer’s patches (**Fig. 4I**). Adherent cells were studied to account for Wnt5A linked bacterial commensalism mostly in connection with dendritic cells, macrophages and macrophage like phagocytes of the Peyer’s patches and to evaluate phagocyte-T cell association in the context of bacterial commensalism, as explained later in the text. The total cells were considered in addition to the adherent cells so as to not miss out on bacteria harboring cells that are weakly adherent.

**Fig. 4.**
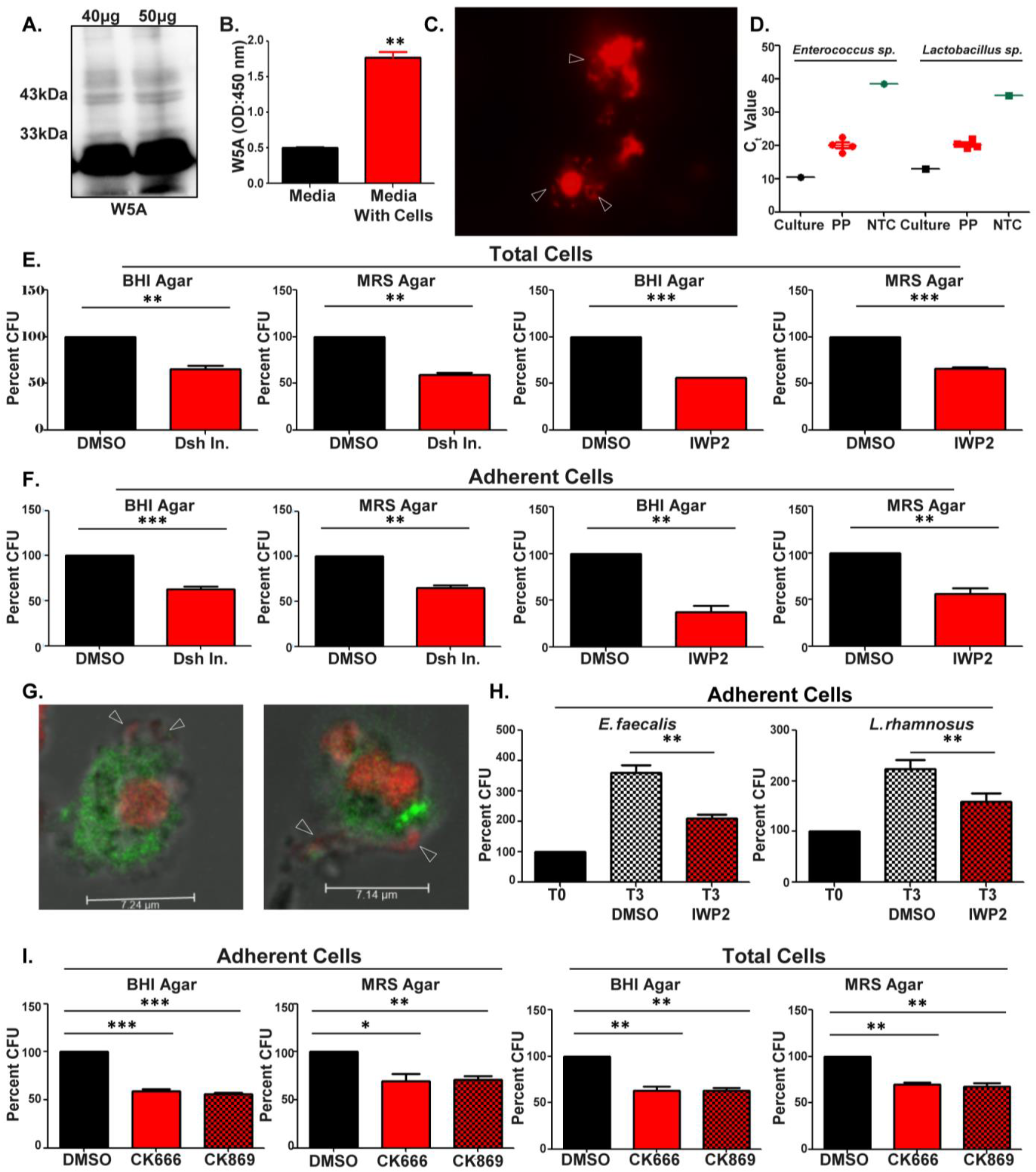
Wnt5A signaling influences commensal bacterial survival in the Peyer’s patches. (A) and (B) Wnt5A expression in cells of Peyer’s patches of BALB/c mice by immunoblot (A) and ELISA (B) (n=2-3). (C) Intracellular bacteria in the adherent cells of Peyer’s patches as observed by fluorescence microscopy after PI staining. (D) Estimation of *Enterococcus sp*. and *Lactobacillus sp*. in Peyer’s patches by qPCR (n=4 mice), NTC indicates No Template Control and ‘Culture’ (*E. faecalis* or *L. rhamnosus*) is used as positive control. (E) and (F) Application of Dsh and Wnt5A production inhibitor IWP2 for 6 hr led to diminution of countable bacterial cfu from total cells (E) and adherent cells (F) of Peyer’s patches (n=3-4). (G) Bacteria (PI: red), denoted by arrow, associated with CD11b (green) phagocytes of Peyer’s patches as observed by confocal microscopy. (H) IWP2 blocked survival of *E. faecalis* and *L. rhamnosus* in adherent cells of Peyer’s patches as observed from cfu 3 hr (T3) post internalization (T0) (n=4). (I) Administration of Arp 2/3 complex inhibitor I (CK666) and Arp 2/3 complex inhibitor II (CK869) for 6 hr to cells of Peyer’s patches led to decrease in endogenous cfu (n=3). Data represented as mean ± SEM and p≤ 0.05 was considered as significant statistically. Significance was represented by * in the following manner: * p≤ 0.05, ** p≤ 0.005, *** p≤ 0.0005.

The influence of Wnt5A on gut bacterial colonization was furthermore reflected in the altered gut bacterial diversity of Wnt5A heterozygous mice, which harbor a single functional copy of the Wnt5A gene, and express less than the wild type level of Wnt5A within the Peyer’s patches (**Fig. 5A and Fig. S6**) and other tissues (19, 36). Since the bacterial composition of fecal matter is used as a signature of gut bacterial colonization (37–39), we characterized the bacterial population of the feces of Wnt5A heterozygous and wild type mice as a measure of bacterial colonization therein. Feces were collected both under normal condition (Day 0) and after antibiotic treatment for 14 days (Day 14) from similarly reared wild type and heterozygous mice (2 from each group) and outsourced for 16S metagenomic sequencing and characterization of bacterial abundance and diversity. Antibiotic was administered specifically to evaluate the potentially protective influence of Wnt5A on gut bacterial abundance/diversity. We found the fecal bacterial colonization of the Wnt5A heterozygous and wild type mice to be markedly different (**Fig. 5B, C** and **D**). Bacterial presence in the feces of the wild type and heterozygous mice both at Day 0 and Day 14 was estimated with respect to a range of different phyla. All phyla present in the different groups (WT: wild type1,2 and HET: heterozygous 1,2) are displayed in the chart and projected as phylum-abundance plot (**5B**). The related Operational Taxonomic Units (OTU: Genus) of the same experimental groups are also projected as alpha diversity using the Shanon, Simpson and Fisher indices (**5C and D**). These results reveal that gut bacterial colonization is more enriched in the wild type mice than in the heterozygous both under normal condition (Day 0) and after antibiotic treatment (Day 14). Additionally, there was considerable difference in the colonization of the *Enterococcus, Lactobacillus, Prevotella* and *Helicobacter* genera within the Peyer’s patches of the Wnt5A heterozygous and wild type mice as validated by qPCR with specific primers (**Fig. 5E**). The Peyer’s patches of the Wnt5A heterozygous mice harbored significantly less *E. faecalis* and *L. rhamnosus* than those of the wild type mice in compliance with the observed dependence of both bacteria on Wnt5A signaling (Fig. 2). Overall, these results indicate that Wnt5A is a regulator of gut bacterial colonization.

**Fig. 5.**
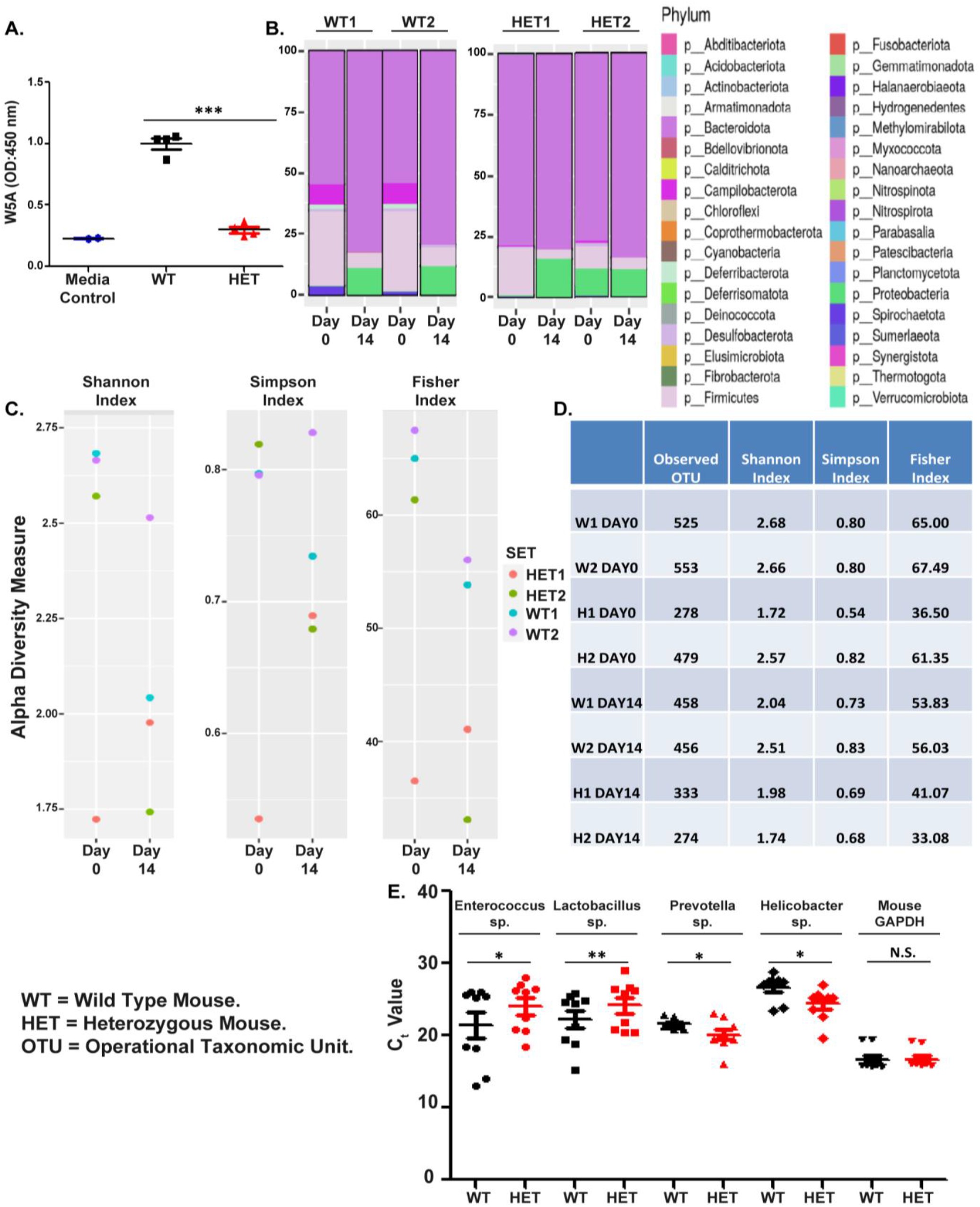
Wnt5A signaling influences colonization diversity of gut commensal bacteria. (A) Lower secretion of Wnt5A from cells of Peyer’s patches of Wnt5A heterozygous mice (lacking a copy of the Wnt5A gene) as compared to its wild type counterpart. (n=4 mice) (B) Difference in fecal bacterial phyla abundance between heterozygous mice 1 and 2 with wild type mice 1 and 2. Observations were made both under normal condition (Day 0) and post Abx treatment for 14 days (Day 14.) (C) Difference in alpha diversity of fecal bacterial population of wild type and heterozygous mice on (Day 0) and after Abx treatment (Day 14) as enumerated using Shannon, Simpson and Fisher indices. (D) Difference in OTU number enumerated from feces of WT (W) and Het (H) mice at Day 0 and Day14 correlating with difference in alpha diversity. (E) qPCR from tissue of Peyer’s patches showing difference in colonization of *Enterococcus, Lactobacillus, Prevotella* and *Helicobacter* in the Wnt5A heterozygous and wild type mice (n=9 mice). Higher C_T_ value indicates lower abundance of bacteria in the particular sample. Data represented as mean ± SEM and p≤ 0.05 was considered as significant statistically. Significance was represented by * in the following manner: * p≤ 0.05, ** p≤ 0.005, *** p≤ 0.0005.

### Regulation of gut commensal bacterial colonization by Wnt5A signaling is linked with T cell activation status

In view of the fact that the gut CD4 T cell repertoire is influenced by the gut resident commensal bacteria (34, 40, 41), we investigated if the altered gut commensal bacterial colonization that correlates with diminution in Wnt5A signaling affects the gut associated T cells. To this end, we studied the activation status of T cells in the Peyer’s patches of Wnt5A wild type and Wnt5A heterozygous mice. Additionally, we analyzed the effect of inhibition of Wnt5A production by IWP-2 on the T cell population of separately isolated Peyer’s patches.

Initially we demonstrated the presence of CD4 T cells in close association with CD11c expressing phagocytes within the mouse gut Peyer’s patches (**Fig. 6A**). CD4 T cell – phagocyte association was similar in both the Wnt5A heterozygous and wild type mice. However, in concurrence with an overall altered gut commensal bacterial abundance and diversity in comparison to that of the wild type controls (Fig. 5), the Wnt5A heterozygous mice harbored CD4 T cells with a distinctly different activation profile within the Peyer’s patches of the gut when compared to the wild type. As demonstrated by FACS in **Fig. 6B** (representative FACS plots) and **6C** (analysis of similar experiments), both the ratio of IL17a+ to Foxp3+ (IL17a/Foxp3) CD4 T cells and the absolute numbers of IL17a+ Foxp3+ double positive CD4 T cells were higher in the Peyer’s patches of Wnt5A heterozygous mice when compared to those of the wild type, indicating alterations in Foxp3 specific regulatory T cell phenotype, IL17a specific activated T cell phenotype and activated vs. regulatory T cell differentiation (33, 34, 42–44) within the Peyer’s patches of the Wnt5A heterozygous mice in relation to the wild type mice. A similar trend toward a relative gain of IL17a specific activated T cell phenotype with reference to a regulatory FoxP3 phenotype and an increase in IL17a+FoxP3+ T cell numbers was also evident when cells isolated from Peyer’s patches were treated with the Wnt5A production inhibitor IWP-2 (**Fig. 6D and E**). Decrease in Wnt5A secretion by cells of the Peyer’s patches and concomitant decrease in the countable bacterial colony forming units therein upon administration of IWP-2 is depicted in **Fig. 4E, F and Figure S5**. These results indicate that as Wnt5A signaling shapes the colonization of gut bacterial flora (**Fig. 4 and 5**), it is also conducive to shaping a bacterial flora dependent gut CD4 T cell repertoire that ranges from a regulatory to inflammatory phenotype depending on the Wnt5A signal dosage. Such a concept is in compliance with the presence of CD4 T cells in close association with the phagocytic cells of the Peyer’s patches (Fig. 6A) and the existence of gut flora specific CD4 T cell repertoire (43–45).The notable correlation of altered MHC II surface expression in cells of the Peyer’s patches with IWP2 mediated reduction in bacterial numbers and alteration in CD4 T cell profile (**Fig. 6F and G**) further suggests that Wnt5A signaling regulates gut CD4 T cell activation status at least partly through MHC II mediated presentation of gut bacterial antigens. Although other investigators have discussed MHC II restricted presentation of bacterial antigens (33), the underlying molecular mechanism of such antigen presentation has remained elusive.

**Fig. 6.**
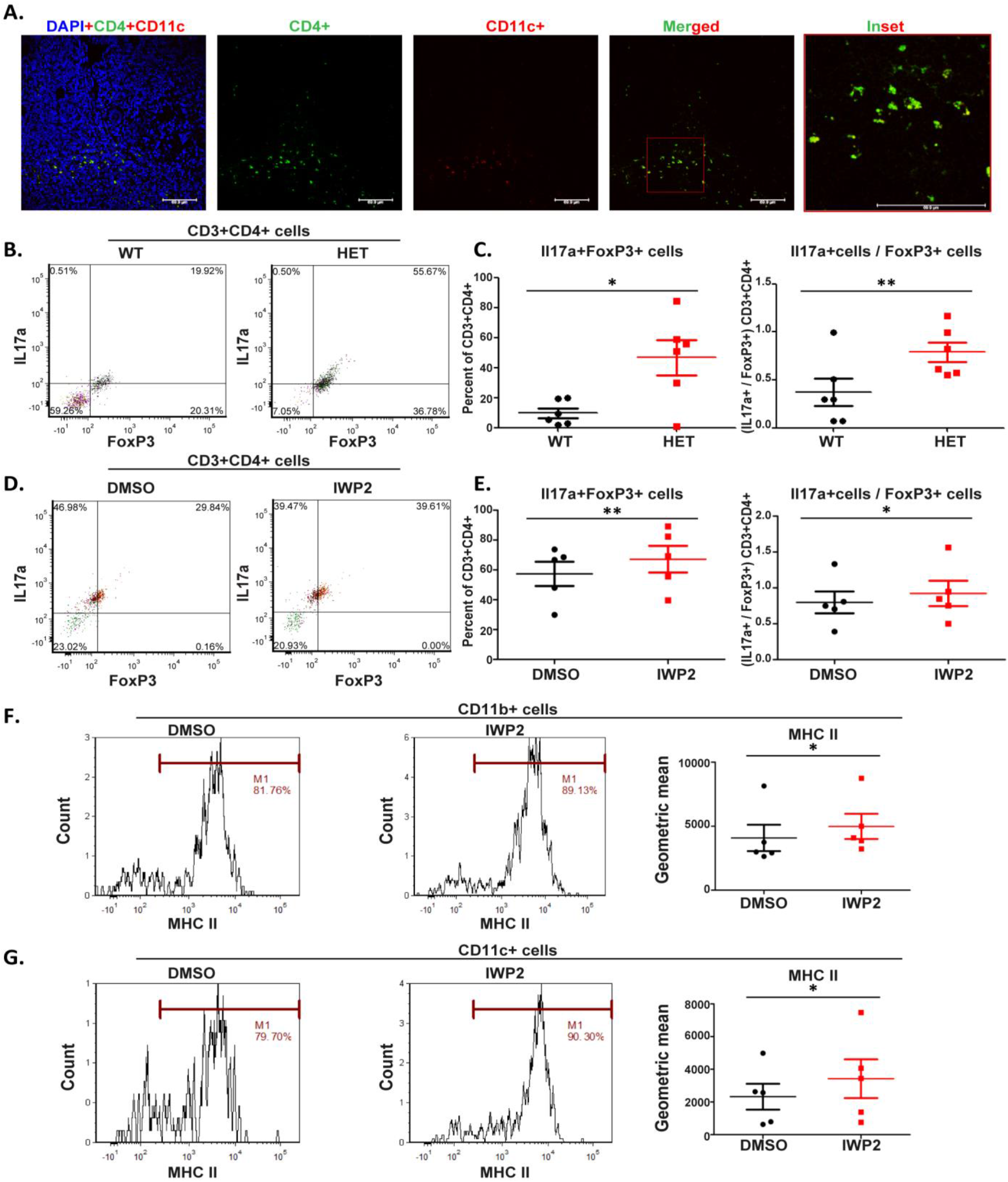
Adaptation of gut commensal bacterial colonization to changes in Wnt5A is linked with altered T cell activation profiles. (A) Colocalisation of CD4+ T cells (green) and CD11c+ phagocytes (red) in Peyer’s patch tissue visualized by confocal microscopy. Inset shows colocalisation (yellow) of both cell types (n=4 mice). (B) FACS representing the difference between IL17a and FoxP3 expressing CD4+T cell population in Peyer’s patches of wild type and heterozygous mice. (C) Plot showing higher IL17a/FoxP3 ratio, as well as higher number of IL17a+FoxP3+ CD4 T cells in Peyer’s patches of heterozygous mice lacking a functional copy of Wnt5A gene as opposed to wild type (n=6 mice). (D) FACS representing the difference in IL17a+ and FoxP3+ expressing CD4 T cell population in cells of Peyer’s patches where Wnt5A signaling was down regulated by IWP2 treatment for 6 hr. (E) Plot showing IWP2 treatment increased IL17a/FoxP3 ratio and number of IL17a+FoxP3+ CD4 T cells in the total cell population of Peyer’s patches (n=5). (F) Higher MHC II surface expression in both CD11b+ and (G) CD11c+ phagocytes of Peyer’s patches on inhibition of Wnt5A signaling by IWP2 (n=5). Data represented as mean ± SEM and p≤ 0.05 was considered as significant statistically. Significance was represented by * in the following manner: * p≤ 0.05, ** p≤ 0.005, *** p≤ 0.0005.

## Discussion

The requirement of gut commensal bacteria for the maintenance of gut immune homeostasis has been reported extensively (20–22, 46, 47), but the molecular basis of interconnection between bacterial commensalism and immune homeostasis is not clearly deciphered. Although one report documents Wnt expression to be associated with gut bacterial colonization (25), the interrelation between steady state Wnt signaling, gut commensal bacterial survival and gut immune homeostasis have not been elucidated in detail. In this study we explained the importance of Wnt5A signaling in the safeguarding of certain commensal bacteria within phagocytes in the steady state. Furthermore, we have described that Wnt5A mediated regulation of commensal bacterial colonization in the murine gut is involved in shaping the gut CD4 T cell repertoire.

Here we demonstrated that a Wnt5A – Actin axis promotes the survival of some common commensal bacteria such as *E* .*faecalis* and *L. rhamnosus* using the macrophage as a model phagocyte. Administration of Wnt5A signaling and actin assembly inhibitors to bacteria harboring macrophages significantly diminishes the bacterial colony forming units therein. Both Wnt5A signaling and the macrophage resident bacteria enrich a self-sustaining circuit that favors Wnt5A production, maintenance of actin assembly and bacterial survival (**Fig. 1 – 3, Fig. S3**). We have also demonstrated the presence of intracellular bacteria within the phagocytes of the gut associated Peyer’s patches. That Wnt5A signaling supports the survival of certain commensal bacteria including *E. faecalis* and *L. rhamnosus* within the Peyer’s patches is evident from the significant reduction in the countable bacterial cfu following treatment of the cells of the Peyer’s patches with either IWP-2, which inhibits Wnt5A production (48, 49), or a Dsh specific inhibitor, which inhibits Wnt5A signaling (27). Similar reduction in the countable bacterial cfu with the application of the actin assembly (Arp2/3) inhibitors suggests that Wnt5A mediated actin modulation contributes to bacterial colonization (**Fig. 4**). The regulatory role of Wnt5A in gut bacterial colonization is further corroborated by the notable differences in both fecal and Peyer’s patch associated bacterial diversity/abundance between the Wnt5A heterozygous mice that lack a functional copy of the Wnt5A gene and their wild type counterparts. The increased prevalence of *Enterococcus* and *Lactobacillus* in the Peyers’ patches of the Wnt5A wild type mice as compared to the heterozygous, and the opposite scenario in case of *Prevotella* and *Helicobacter* are quite likely due to variation in the interactions of the existing bacteria with host Wnt5A signaling on account of difference in their pathogenic potential (**Fig. 5**). In harmony with being conducive to colonization of certain bacterial species in the gut, Wnt5A signaling also supports the maintenance of a gut specific CD4 T cell repertoire. This is evident from the observed change in CD4 T cell activation that correlates with Wnt5A mediated regulation of gut bacterial colonization (**Fig. 6**). Significant change in the profile of the CD8 T cells of the gut was not observed (**Fig. S9**).

In light of the existing literature describing commensal bacteria specific CD4 T cells (22, 33), it is quite possible that the Wnt5A-Actin axis of the phagocytes of the Peyer’s patches orchestrates presentation of resident bacteria derived epitopes as self-antigens to CD4 T cells thereby regulating their activation. This concept is in compliance with the presence of bacteria harboring phagocytes and their co-existence with CD4 T cells within the Peyer’s patches (**Fig. 6A)**. Depending on the extent and affinity of MHCII-bacterial peptide –CD4 T cell receptor interactions, the CD4 T cells could be accordingly sensitized at the transcriptional level leading to the development of an activated (for example, IL17a secreting) or regulatory (for example, Foxp3 expressing) phenotype (22, 50). Thus, the Wnt5A-Actin axis of phagocytes could play a major role in shaping the gut steady state CD4 T cell repertoire in the context of the incumbent bacteria.

Taken together, the data presented in this manuscript highlight that Wnt5A signaling contributes to the maintenance of gut CD4 T cell homeostasis by sustaining a steady commensal bacterial colonization. Although we have focused mostly on the Wnt5A-Actin axis of macrophages or macrophage like cells in relation to the interconnection between gut bacterial commensalism and gut CD4 T cell homeostasis, the involvement of other immune sensors therein certainly cannot be ruled out. In this connection investigation into the involvement of TLR (51–53), NOD2 (54–56) or other autophagy inducing Pattern Recognition Receptors, which have not been covered in this study should be of considerable interest. Nevertheless, this study upholds the requirement of Wnt5A signaling as a crucial regulatory factor of gut immune homeostasis, lack of which may result in defective host-commensal mutualism leading to dysbiosis and the development of autoimmune disorders (24, 56).

## Materials and Methods

### Cells and Reagents

#### Cells

RAW264.7 cell line was obtained from ATCC (ATCC©TIB71™). Peritoneal macrophages and cells of Peyer’s patches were harvested and cultured following previously published protocols (17, 19, 57–59). Cells were grown at 37°C with 5% CO_2_ in DMEM High Glucose containing 10% FBS, 2 mM L-glutamine, 100 U/ml penicillin and 100 mg/ml streptomycin. *E. faecalis* was isolated from the cecum of BALB/c mouse. *L. rhamnosus* and *S. enterica* were purchased from MTCC, Chandigarh, India (MTCC 1408^T^, MTCC 3224).

#### Reagents

List of reagents used is provided in Supplementary Information Table S1.

### Isolation and characterization of commensal bacteria

For isolation of commensal bacteria from cecum of BALB/c mouse, the cecum was perforated with a needle. Subsequent inoculation of BHI broth with the needle and overnight incubation produced enough bacteria, which were then plated in BHI agar. After 12 hr colonies were plucked from which pure cultures were generated. Accordingly, bacterial stocks were made for 16S sequencing and used for experiments thereafter.

### Isolation and culture of cells from Peyer’s patches

Small Intestine was harvested from BALB/c or Wnt5A wild type (+/+) and Heterozygous (+/-) mice as per need. Intestine was kept moist and flushed with sterile PBS to remove fecal material. Peyer’s patches were identified opposite to the mesenchymal end of the Intestine and removed as sections of 1 mm length. Peyer’s patches were then shaken at 200 RPM under 37° C in RPMI 1640 media without antibiotic supplemented with 10% FBS and 10 mM EDTA for 20 min. Media was then decanted and replenished with RPMI 1640 (+10% FBS, -Antibiotic) with added Collagenase D and DNAse I (AM2222) following which the Peyer’s patches were shaken for an additional 15 min. After dispersion of the tissue and its passage through 40 µM cell strainer the dispersed mix with some isolated cells was again shaken at 200 RPM for 10 min. The mix was then passed through another 40 µm strainer to get single cell population. These cells were used for Flow Cytometry or cultured with RPMI 1640 containing 10% FBS and 2 mM L-glutamine, either with or without antibiotic (100 U/ml penicillin and 100 mg/ml streptomycin) as per experimental need.

### Bacterial Uptake Assay

RAW264.7 cells or mouse peritoneal macrophages were pre-treated with recombinant Wnt5A (rWnt5A) (50ng/ml, dissolved in PBS) for 6 hr, following which the cells were infected with either *E. faecalis* or *L. rhamnosus* at MOI 10 and 20 respectively for 2 hr in DMEM High Glucose media without antibiotic and FBS. Cells were then washed with PBS, lysed in sterile water and plated in either BHI (*E. faecalis*) or MRS (*L. rhamnosus*) Agar. Cfu were counted after 14 hr (BHI) or 36 hr (MRS) of incubation under 37° C with (MRS) or without (BHI) 5% CO_2_.

### Bacterial Survival/Killing Assay

RAW264.7 cells or peritoneal macrophages were pre-treated with recombinant Wnt5A (rWnt5A) or PBS for 6 hr and infected with *E. faecalis* (MOI 10) or *L. rhamnosus* (MOI 20) for 1 hr After washing off the external bacteria with PBS, the infected cells were either harvested to estimate bacterial uptake (time point T0) or incubated with DMEM High Glucose containing 10% FBS, 2 mM L-glutamine, devoid of antibiotic, to be harvested at 1 hr intervals (T1-T4) for assessing killing or survival of the internalized bacteria at the different time points. As explained earlier, each batch of the harvested cells were lysed and plated in BHI or MRS agar. Cfu calculation was done after incubation for 14 hr (BHI) or 36 hr (MRS) under 37° C with (MRS) or without (BHI) 5% CO_2_.

For observing the effect of Dsh (15 µm), Rac-1 (15 µm) or actin polymerization inhibitors CK 666 & CK 869 (20 µm each) on bacterial survival, infected cells were incubated in antibiotic free DMEM +10% FBS, in the presence of the desired inhibitors for 3 hr (T3). After harvesting at T3, the cells were lysed and plated in MRS or BHI agar for estimating cfu. In case of Fz5 antiserum application, the same experiments were performed with the only exception that cells were harvested 3 and 6 hr (T3, T6) post infection removal.

Similar bacterial survival assays were performed with cells obtained from Peyer’s patches using IWP2 (0.05 µm) and Dsh (15 µm) inhibitors. Cells were first incubated for 12 hr in RPMI 1640 supplemented with 10% FBS, 2 mM L-glutamine, 100 U/ml penicillin and 100 mg/ml streptomycin under normal tissue culture conditions to minimize cell associated bacteria. Subsequently, the treated cells were infected with *E. faecalis* (MOI 10) or *L. rhamnosus* (MOI 20) for 1 hr, and after removal of the external bacteria were incubated in antibiotic free RPMI 1640 containing 10% FBS and any one of the desired inhibitors for 3 hr cfu calculation was done in same manner as previously explained.

Survival of endogenous bacteria was estimated after isolation of cells from Peyer’s patches. Cells were grown for 6 hr at 37°C with 5% CO_2_ in RPMI 1640 supplemented with 10% FBS, 2 mM L-glutamine along with Wnt signaling inhibitors namely Dsh inhibitor(15 µM), IWP2 (0.05 µM) or actin assembly inhibitors CK666 (20 µM), CK869 (20 µM) or DMSO as vehicle control and devoid of any antibiotic. After incubation cells were harvested, lysed in sterile water and plated in BHI and MRS agar. Further incubations and cfu calculations were done following earlier mentioned procedures.

### Transfection

RAW264.7 macrophages were first plated for 6-8 hr in six well tissue culture plates (approx. 2×10^6^ cells per well) and incubated at 37°C with 5% CO_2_ for infection and subsequent transfection. Cells were infected with either *E. faecalis* or *L. rhamnosus* at MOI 10 or 20 for 1hr in DMEM devoid of FBS and Pen-Strep. After removal of the external bacteria, the infected cells were incubated for 2 hr in DMEM supplemented with 10% FBS, 2 mM L-glutamine, 100 U/ml penicillin and 100 mg/ml streptomycin to ensure complete removal of all the remaining extracellular bacteria. Subsequently, the medium was replaced by transfection mix as follows. Wnt5A siRNA (50 µM) or Wnt3A (100 µM) siRNA or non-targeting (control) siRNA(100 µM) separately mixed with 5 µl with Lipofectamine RNAiMAx reagent in 300 µl of antibiotic free serum free OPTI-MEM medium was incubated for 30 min, diluted to 25nM in 700 µl antibiotic free DMEM with 2% FBS, and added to cells in each well. After 24 hr the culture was replaced with DMEM (+2 mM L-glutamine, +10% FBS, +Pen-Strep) and incubated further for 36 hr Cells were then washed, lysed and plated in BHI or MRS agar as previously explained. Cfu was calculated controlled to cell number.

### Western Blotting

Cells were harvested, and lysis was done with lysis buffer for 15 min at 4°C after which centrifugation was done for 10 min at 16,000 g in 4°C. Approximately 40 µg of the lysate was loaded into each well and SDS-PAGE was run. Proteins from the gel were transferred to PVDF membrane. Blocking was done with 5% BSA for 2 hr following which the membrane was incubated overnight with primary antibody at 4°C. Incubation with appropriate HRP conjugated secondary antibody was continued for 2 hr at room temperature with mild shaking. Finally visualization of the membrane was done by chemiluminescent substrate using Chemi documentation system of Invitrogen and iBright FL-1500 and Azure Biosystems, Model-C400. Band intensities were calculated utilizing GelQuant.Net.

### Confocal/Fluorescence Microscopy

#### PI (Propidium Iodide) staining

For identification of intracellular bacteria propidium Iodide staining was performed with minor modification of previously published protocol (19). Peritoneal macrophages plated in 4 chambered glass slides were pretreated with either rWnt5A or PBS before infection with *E. faecalis* (MOI 10) or *L. rhamnosus* (MOI 20) for 2 hr, following which the cells were fixed with 4% paraformaldehyde for 10 min. Fixed cells were stained with 30 µM PI in PBS containing 0.1% Triton X for 15 min at room temperature. After washing of excess of the stain with PBS mounting was done with 60% glycerol. The cells were observed under fluorescence microscope (Leica DMI3000 B) at 100X objective and 1X zoom. For cells isolated from Peyer’s patches plating was done in Poly-L-Lysine coated chambered slides. Cell were plated in DMEM High glucose (+10% FBS,- PenStrep) for 3 hr and then were treated similarly a in the previous cases. For CD11b staining cells were stained with the primary antibody overnight at 4°C followed by incubation with flurophore tagged secondary antibody for 2 hr PI staining was done thereafter following similar protocol as previously mentioned. Cells were visualized under Leica TCS-SP8) at 60X magnification and 4X Zoom

#### Phalloidin Staining

Phalloidin staining was performed for visualizing actin assembly.RAW264.7 cells were plated in 4-chambered slides and either infected with *E. faecalis* (MOI 10) or *L. rhamnosus* (MOI 20) were incubated for 2 hr under normal tissue culture conditions following which cells were fixed with 4% paraformaldehyde for 10 min. Phalloidin (Alexa Fluor 555) and DAPI diluted to 1:2000 and 1:4000 ratios respectively by 2.5% BSA in PBST(0.1% Tween-20) was added to the fixed cells and incubation was continued for 15 min. Cells were then washed 3 times and mounted with 60% glycerol before observation by confocal microscope Zeiss LSM 980 at 60X magnification and 2X zoom.. Transfected cells were treated similarly for microscopy and visalized using either Zeiss LSM 980 or (Leica TCS-SP8) at 60X magnification and 2X zoom.

#### Antibody staining of paraffin section of Peyer’s patches

Peyer’s patches were isolated and fixed overnight in 10% buffered formalin. After fixation tissues were embedded in paraffin and 3 µm sections were made by microtome. Sections were then deparaffinized and rehydrated using following treatment of slides using Xylene 2 × 3 min, Xylene 1:1 with 100% ethanol for 3 min, 100% ethanol 2 × 3 min, 95% ethanol for 3 min, 70% ethanol for 3 min, 50% ethanol for 3 min and lastly rinsed with cold tap water. Heat induced epitope retrieval was done using Tris-EDTA buffer (10 mM Tris base, 1 mM EDTA, 0.05% Tween 20, pH 9.0). Antibody staining was done using 1% BSA in TBS. Tissue with primary antibody was incubated overnight at 4°C followed by incubation with alexa fluor 488 conjugated secondary antibody for 2 hr at room temperature. For flurophore conjugated primary antibodies staining was done for 2 hr at room temperature. Tissue sections were visualized under Leica TCS-SP8) at 40X magnification and 0.75X Zoom

### Flow Cytometry

For flow cytometry, cells isolated from Peyer’s patches were plated, treated as per need and allowed to settle. Treatment with Brefeldin A was given to cells at concentration of 3 µg/ml for 4 hr for checking intracellular cytokine or transcription factor. After washing with PBS, the cells were fixed with 1% paraformaldehyde and permeabilized with PBST (0.5% tween 20). Subsequently antibody staining was done with the desired antibodies in 1% BSA in PBST for 1 hr at 4°C.For observation of phagocytes and surface MHC II expression live cells post treatment and without Brefeldin A exposure were stained with antibodies in a similar manner as above before fixation with 1% paraformaldehyde. Flow cytometry was done by BD.LSR Fortessa Cell analyser and analysis was done via FCS Express 5 software from BD (Beckton Dickinson).

### Analysis of fecal bacterial diversity

Fecal pellets were collected from Wnt5A wild type and Wnt5A heterozygous mice initially without any treatment. After that all mice were treated with Abx cocktail (4 mg/mL kanamycin,0.35 mg/mL gentamicin, 8,500 U/mL colistin, 2.15 mg/mL metronidazole and 0.45 mg/mL vancomycin) (60) for 14 days and fecal pellets were again harvested. All pellets were sent for 16S sequencing and data analysis to LCGC Life Sciences/WIPRO. Quality assessments of the isolated genetic materials were done by D1000 ScreenTape assay of TapeStation systems. Sizing range was 35-1000 bp with 15% resolution between 35-300bp and 10% between 300-1000 bp. Trimming of adapters and poor read sequences were done by Trim Galore. Quality profiling and error estimation of sequence reads was conducted utilizing R package dada2. Kraken 2 and Bracken tools were used to align the filtered reads in relation to 16S rRNA annotations from Silva v138 database. Phylogenetic analysis was done generating plots and statistics for taxonomic classification by R package phyloseq. Visualization of interactive hierarchial chart representing taxonomic classification was done using Krona tool.

### RNA isolation for qPCR

Peyer’s patches were excised after cleaning with PBS and homogenized in 600 µl Trizol 2 – 3 times for 2 min on ice, following which120 µl of chloroform was added to each homogenate and mixed well. After centrifugation at 16,000 g for 12 min the upper transparent phase was collected, to which 300 µl of isopropanol was added. After mixing and incubation on ice for 30 min, centrifugation was performed at 16,000 g for 12 min. After decanting supernatant, the pellet was washed with 70% ethanol and left to air dry. Dried RNA pellet was dissolved in DEPC treated water by heating at 50°C. Gene specific cDNA was made with standard RT kit (Biobharati Life Sciences) using reverse primers specific to 16S genes of the bacterial strains we wanted to observe. qPCR of cDNA samples were run in CFXOpus96 of Biorad. Amplification of 16S region of *Enterococcus sp*., *Lactobacillus sp*., *Prevotella sp*., *Helicobater sp*. and mouse GAPDH was visualized using SYBR green dye. C_t_ values were plotted using Graphpad Prism 5 for statistical analysis. List of primers used is provided in table in supplementary information Table S2.

### Isolation of bacterial DNA for PCR

Bacterial genomic DNA was isolated from cells of Peyer’s patches using Qiagen DNeasy Blood and Tissue Kit with slight modifications of previously published protocol (61) for lysis of bacteria. Approximately 4 × 10^8^ cells were resuspended in 180 µl of Enzymatic lysis buffer (20 mM Tris-Cl, pH 8.0, 2 mM Sodium EDTA, 1.2% Triton X-100 and 20 mg/ml lysozyme freshly added before use) and incubated at 37°C for 1 hr, following which 20 µl Proteinase K (20 mg/ml) was added to the mix and further incubated overnight at 56°C. Further 200 µl of buffer AL (Lysis buffer) was added and incubated for 30 min at 56°C, after which 200 µl of 100% Ethanol was added and mixed thoroughly. The mix was loaded onto DNeasy mini spin column and centrifuged for 1 min, at 7,200 g. Flowthrough was discarded and 500 µl of wash buffer AW1 was added and centrifugation was performed as before. Further discarding the flowthrough, 500 µl of wash buffer AW2 was added followed by centrifugation at 14,000 g for 4 min. DNA elution was done using 50 µl of elution buffer by centrifugation at 7,200 g for 1 min. Eluted Genomic DNA was used in PCR as per requirements.List of PCR primers are in upplementary information Table S2.

### ELISA

Indirect ELISA was performed to analyze the level of Wnt5A or Wnt3A secreted from cells. In case of RAW264.7 cell culture (infected or un-infected), the harvested culture media was centrifuged at 7,200 g for 3 min to remove extracellular bacteria and coated in 96 well ELISA plate for overnight incubation at 4°C. Following decanting and washing with PBST (0.05% Tween 20) 3 times, blocking was done with 1% BSA in PBS for 2 hr Plates were then incubated overnight at 4°C with primary antibodies against Wnt5A (1:1000 dilution) or Wnt3A (1:2000 dilution). After further washing with PBST 3 times HRP conjugated anti-Rat secondary antibody was added at 1:4000 dilution and incubation was continued for 2 hr After washing 3 times with PBST the HRP substrate TMB was added to each well. Stop solution (2N H_2_SO_4_) was added after 15 min to stop the color reaction. Reading was taken at 450nm in ELISA plate reader. For checking the level of secretion of Wnt5A from the Peyer’s patches of Wnt5A wild type and heterozygous mice, the culture medium was harvested from cells of the respective Peyer’s patches and analyzed in the same manner.

### Statistical Analysis

Statistical analysis of data were done utilizing Paired or Unpaired Student t test as per need in Graphpad Prism 5 software. Graphs and line diagrams are represented as mean ± SEM and p≤ 0.05 was considered as significant statistically. Significance was represented by * in the following manner: * p≤ 0.05, ** p≤ 0.005, *** p≤ 0.0005.

## Ethics Statement

All animal experiments were conducted with approval of Animal Ethics committee of CSIR-IICB as per meeting IICB/AEC/Meeting/Sep/2019/1 held on 19/09/2019.

## Acknowledgments

The authors acknowledge Tresa Rani Sarraf and Tanmoy Dalui for assistance in FACS, Shounak Bhattacharya and Vivek Chander for assistance in confocal microscopy, CSIR-IICB Central Instrument Facility for instrument support. We acknowledge Dr. A. Konar and CSIR-IICB animal house facility for assistance with animal breeding and maintenance. We acknowledge Megha Mallick for her assistance in identification of commensal bacteria and Chandan Bhattacharya for histological tissue processing.

