## Supplemental Information for "Wnt5A Signaling Regulates Gut Bacterial Survival and T cell Homeostasis"

**Table S1:** List of reagents used in the study

| Reagent or Resource | Source | Ref. No. |
|---|---|---|
| DMEM High Glucose | Life Technologies (Thermofisher Scientific, USA) | 12800-017 |
| RPMI 1640 | Life Technologies (Thermofisher Scientific, USA) | 31800-022 |
| HI FBS | Life Technologies (Thermofisher Scientific, USA) | 10082147 |
| Penicillin-Streptomycin | Life Technologies (Thermofisher Scientific, USA) | 15140122 |
| L-glutamine | Life Technologies (Thermofisher Scientific, USA) | 25030081 |
| 1X Trypsin EDTA | Life Technologies (Thermofisher Scientific, USA) | 25200-072 |
| Trizol reagent | Life Technologies (Thermofisher Scientific, USA) | 15596018 |
| DAPI | Life Technologies (Thermofisher Scientific, USA) | D1306 |
| Alexa Fluor 555 Phalloidin | Life Technologies (Thermofisher Scientific, USA) | A34055 |
| Brain Heart Infusion broth (BHI) | HiMedia Laboratories,India | M210 |
| MRS broth | HiMedia Laboratories,India | M369 |
| Agar agar | HiMedia Laboratories,India | GRM666 |
| Anti-β actin antibody | Santacruz Biotechnology, USA | SC-47778 |
| IWP2 | Santacruz Biotechnology, USA | SC-252928 |
| Anti-mouse Wnt5A antibody | R&D systems, USA | MAB-645 |
| Anti-mouse Wnt3A antibody | R&D systems, USA | MAB-1324 |
| Anti-rat HRP conjugated 2° antibody | R&D systems, USA | HAF005 |
| Anti-mouse CD4 | R&D systems, USA | MAB-554 |
| Anti-mouse HRP conjugated 2° antibody | Sigma-Aldrich, USA | A9044 |
| Collagenase D | Sigma-Aldrich, USA | 11088866001 |
| Kanamycin Sulphate | Sigma-Aldrich, USA | K4000 |
| Gentamycin | Sigma-Aldrich, USA | G1914 |
| Colistin | Sigma-Aldrich, USA | C4461 |
| Metronidazole | Sigma-Aldrich, USA | M1547 |
| Vancomycin | Sigma-Aldrich, USA | V2002 |
| Lysozyme | Sigma-Aldrich, USA | L6876 |
| Glycerol | Sigma-Aldrich, USA | G5516 |
| NaCl | Sigma-Aldrich, USA | S5886 |
| NP-40 | Sigma-Aldrich, USA | 492018 |
| Triton X-100 | Sigma-Aldrich, USA | 11332481001 |
| Anti-Fz5 antiserum, raised in rabbit | Biobharati Life Sciences, India | |
| CDNA synthesis kit | Biobharati Life Sciences, India | BB-E0043 |
| Taq Polymerase | Biobharati Life Sciences, India | BB-E0010 |
| DMSO | MP- Biomedicals, USA | 196055 |
| rWnt5A | Merck, Germany | GF146 |
| Rac1 Inhibitor | Merck, Germany | NSC23766 |
| Disheveled (Dsh-PDZ domain) Inhibitor | Merck, Germany | CAS294891-81-9 |

| Reagent or Resource | Source | Ref. No. |
|---|---|---|
| Arp 2/3 complex Inhibitor I | Merck, Germany | CK-666 |
| Arp 2/3 complex Inhibitor II | Merck, Germany | CK-869 |
| Tris Base | Merck, Germany | 648310 |
| $Na_3VO_4$ | Merck, Germany | D00152519 |
| TMB solution | Merck, Germany | CL07-1000MLCN |
| PVDF membrane | Millipore, USA | IPVH00010 |
| Lumintaclassico chemiluminescent substrate | Millipore, USA | WBLUC0500 |
| $MgCl_2$ | Millipore, USA | 60583305001046 |
| Tween-20 | Millipore, USA | 655205 |
| β-mercaptoethanol | Millipore, USA | 8057400250 |
| NaF | Millipore, USA | 61773705001730 |
| SMART pool mouse Wnt5A siRNA | Dharmacon, USA | L-065884-01 |
| SMART pool mouse Wnt3A siRNA | Dharmacon, USA | 046386-00-0005 |
| Non targeting pooled Control siRNA | Dharmacon, USA | D-001810-05 |
| V500 syrian hamster anti-mouse CD3 | BD Biosciences, USA | 560771 |
| PerCP-Cy 5.5 rat anti-mouse CD4 | BD Biosciences, USA | 550954 |
| BV605 rat anti-mouse CD8 | BD Biosciences, USA | 563152 |
| Alexa fluor 488 rat anti-mouse IL17A | BD Biosciences, USA | 560221 |
| Alexa fluor 647 rat anti-mouse FoxP3 | BD Biosciences, USA | 560401 |
| FITC rat anti-mouse CD11b | BD Biosciences, USA | 553310 |
| PE rat anti-mouse CD11b | BD Biosciences, USA | 557397 |
| APC hamster anti-mouse CD11c | BD Biosciences, USA | 550281 |
| PE mouse anti-mouse H-2K[d] | BD Biosciences, USA | 553566 |
| PerCP-Cy 5.5 rat anti-mouse I-A/I-E | BD Biosciences, USA | 562383 |
| FITC mouse anti-mouse H-2Kb | BD Biosciences, USA | 553569 |
| Propidium Iodide | BD Biosciences, USA | 556463 |
| Fixable Viability Stain 450 | BD Biosciences, USA | 562247 |

**Table S2**: List of qPCR primers used in the study

| Organism | Primer Name | Primer Sequence (5'-3') | No. of bases |
|---|---|---|---|
| *Enterococcus sp.* | EF_ID_Fw | GCAAGTCGAACGCTTCTTTC | **20** |
| | EF_ID_Rv | GCACCTGTTTCCAAGTGTTATC | **22** |
| *Lactobacillus sp.* | Lac_ID_Fw | GAAGGCTTTCGGGTCGTAAA | **20** |
| | Lac_ID_Rv | CGTGGCTTTCTGGTTGGATA | **20** |
| *Prevotella sp.* | Prev_ID_Fw | GATGCGTCTGATTAGCTTGTTG | **22** |
| | Prev_ID_Rv | CCGTGTCTCAGTTCCAATGT | **20** |
| *Helicobacter sp.* | Hel_ID_Fw | GGAATCACTGGGCGTAAAGA | **20** |
| | Hel_ID_Rv | CCTCTCCCACACTCTAGACTAATA | **24** |
| Mouse | Mo_RNA_GAPDH_Fw | AACAGCAACTCCCACTCTTC | **20** |
| | Mo_RNA_GAPDH_Rv | CCTGTTGCTGTAGCCGTATT | **20** |

**Table S3**: List of PCR primers used in the study

| Organism | Primer Name | Primer Sequence (5'-3') | No. of bases |
|---|---|---|---|
| *Enterococcus sp.* | EF2FP | CAGTTACTAACGTCCTTGTTC | 21 |
| | EF2RP | CGCTTCTTTCCTCCCGAGT | 19 |
| *Lactobacillus sp* | LRFP | GGTGTTGGAGGGTTTCCG | 18 |
| | LRRP | CTAAATGCTGGCAACTAGTC | 20 |

### Supplementary Fig. 1

A.

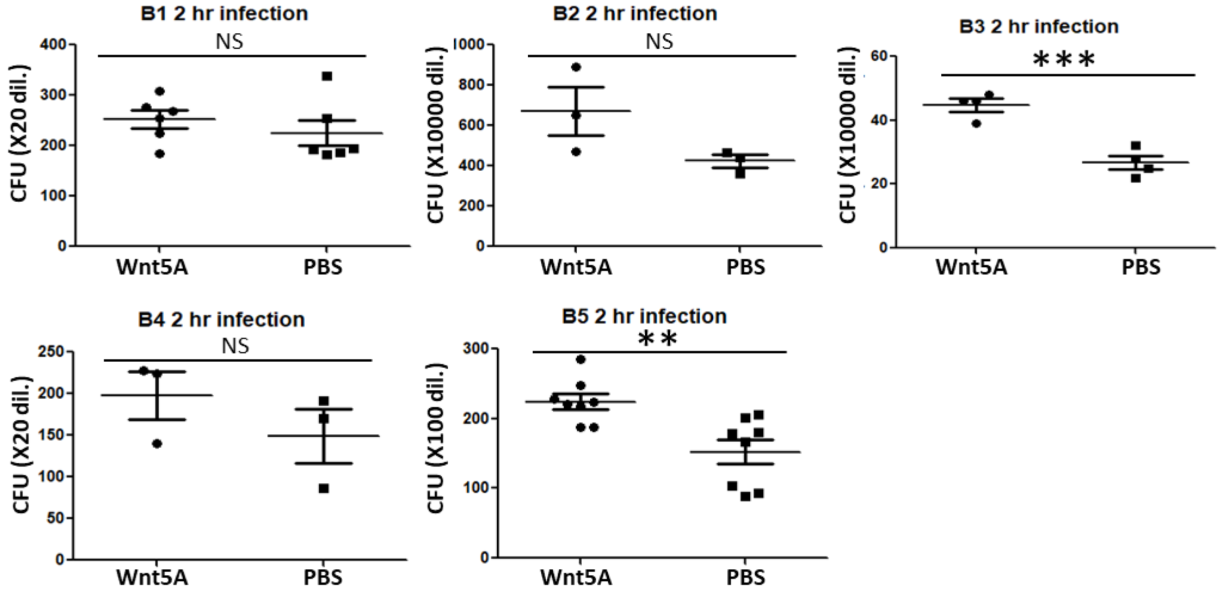


B.

| Description | Query Cover | Per. Ident | Accession |
|---|---|---|---|
| *Enterococcus faecalis* strain ATCC 19433 16S ribosomal RNA, partial sequence | 100% | 99.18% | NR_115765.1 |
| *Enterococcus faecalis* strain LMG 7937 16S ribosomal RNA, partial sequence | 100% | 98.98% | NR_114782.1 |

C.

| Description | Query Cover | Per. Ident | Accession |
|---|---|---|---|
| *Lactobacillus rhamnosus* strain NBRC 3425 16S ribosomal RNA, partial sequence | 99% | 99.67% | NR_113332.1 |
| *Lactobacillus rhamnosus* strain JCM 1136 16S ribosomal RNA, partial sequence | 99% | 99.33% | NR_043408.1 |

**Fig S1: Isolation and Identification of Wnt5A responsive bacterial commensals.** (A) Bacteria isolated from cecum of BALB/c mice were tested for responsiveness towards Wnt5A signaling by estimating changes in internalization by RAW264.7 cells pre-treated with rWnt5A or PBS (vehicle control). Isolates were named B1-B5 for convenience of identification. Isolates B3 and B5 showed highest responsiveness towards Wnt5A signaling.(n=3-8) (B) Isolate B3 was identified as *E. faecalis* by 16S sequencing. (C) Identity of commercially acquired *L. rhamnosus* was also verified by 16S sequencing. “n” represents number of experiments. Data represented as mean ± SEM and p≤ 0.05 was considered as significant statistically. Significance was represented by * in the following manner: * p≤ 0.05, ** p≤ 0.005, *** p≤ 0.0005. “N.S.” denotes non-significant.

**Supplementary Fig. 2**

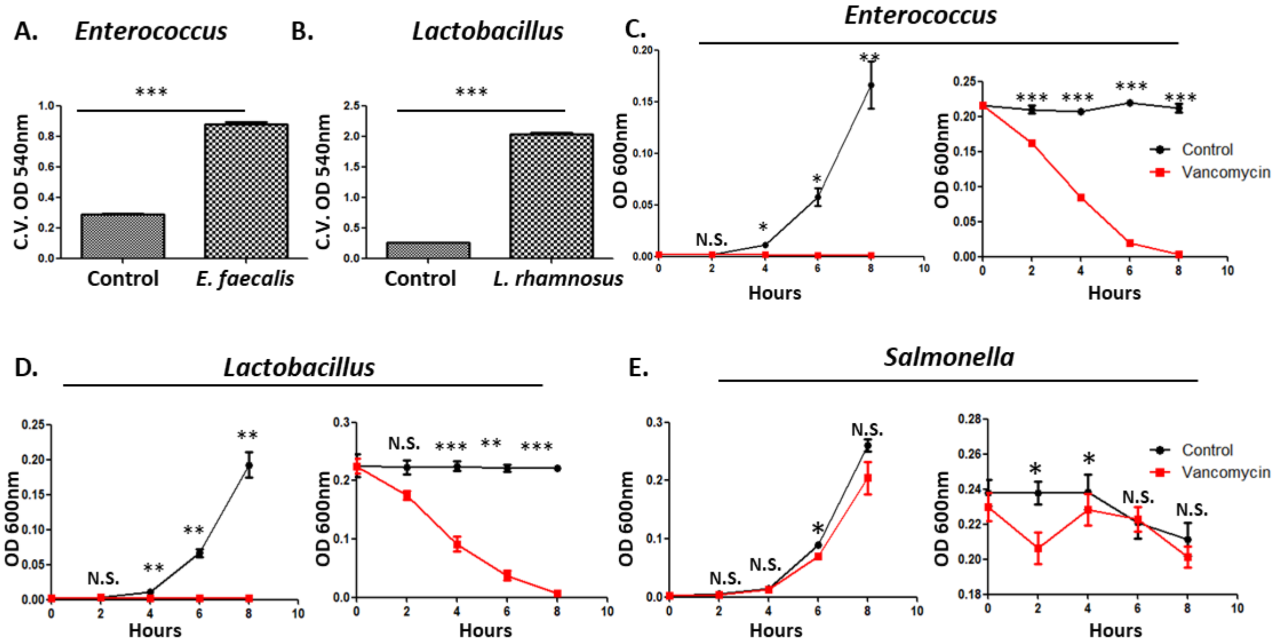


**Fig S2: Characterization of *E. faecalis* and *L. rhamnosus*.** (A) *E. faecalis* and (B) *L. rhamnosus* displayed biofilm forming potential, a common feature reported in gut commensals. Biofilm formation was observed by Crystal Violet (CV) staining (n=3). (C) *E. faecalis* and (D) *L. rhamnosus* lacked pathogenicity as confirmed by lack of vancomycin resistance. Newly inoculated bacterial culture failed to grow and grown culture died out when CDC recommended 6µg/ml vancomycin was added in growth medium and observed with respect to control (without vancomycin) (n=3). (E) *S. enterica* , a well-established gut pathogen displayed Vancomycin resistance (n=3). “n” represents number of experiments. Data represented as mean ± SEM and p≤ 0.05 was considered as significant statistically. Significance was represented by * in the following manner: * p≤ 0.05, ** p≤ 0.005, *** p≤ 0.0005.

#### Supplementary Fig. 3

A.

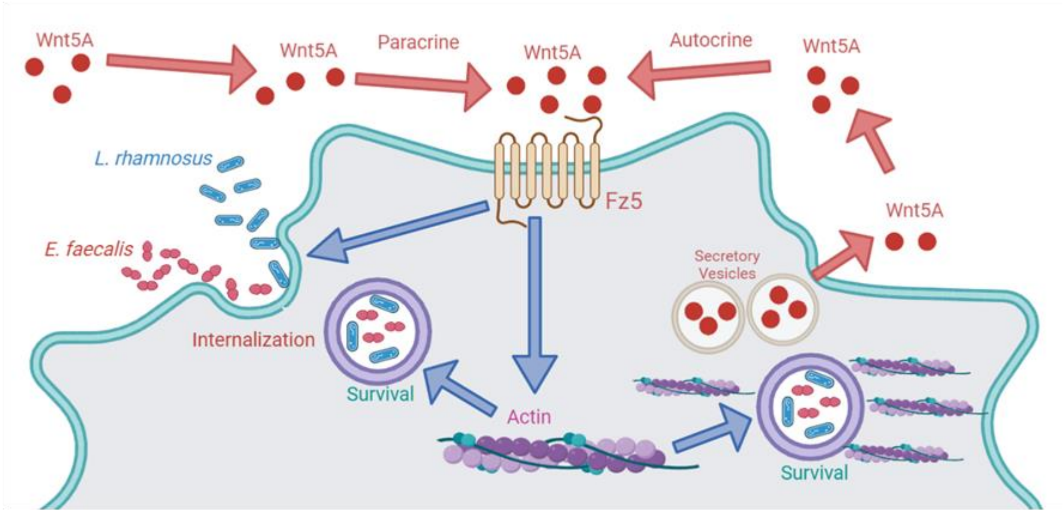


B.

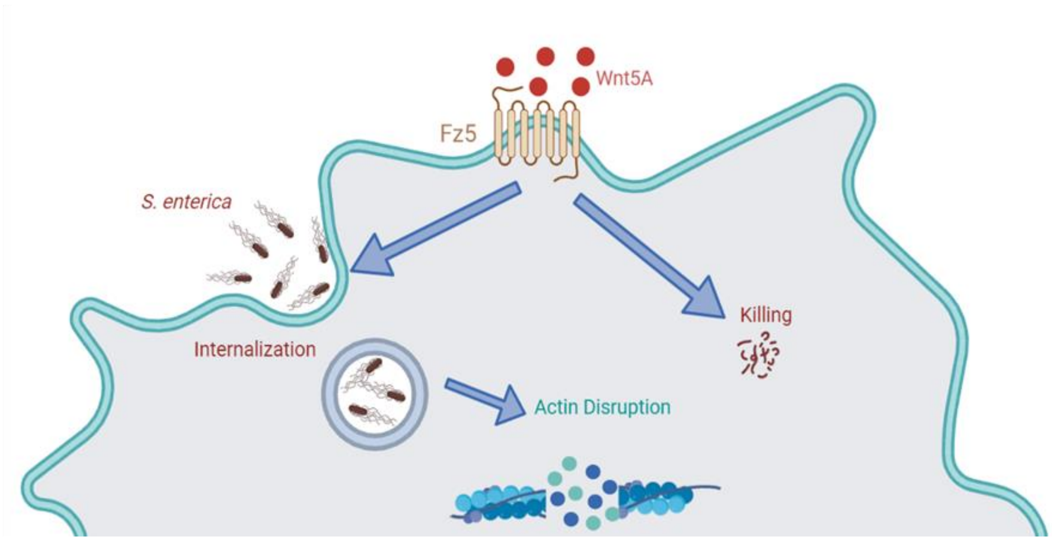


**Fig S3: Model depicting Wnt5A-Actin axis mediated survival of bacterial commensals through a self perpetuating circuit .** (A) Wnt5A aids the internalization of bacterial commensals which are in harmony with the Wnt5A-Actin axis. While the survival of the internalized commensals is bolstered by the host, commensal internalization enhances Wnt5A secretion, leading to a self sustaining Wnt5A-Actin-Commensal loop. (B) Wnt5A upregulates internalization of the pathogen *S. enterica* but the pathogen disrupts the host actin machinery, causing Wnt5A signaling to antagonize its survival.

### Supplementary Fig. 4

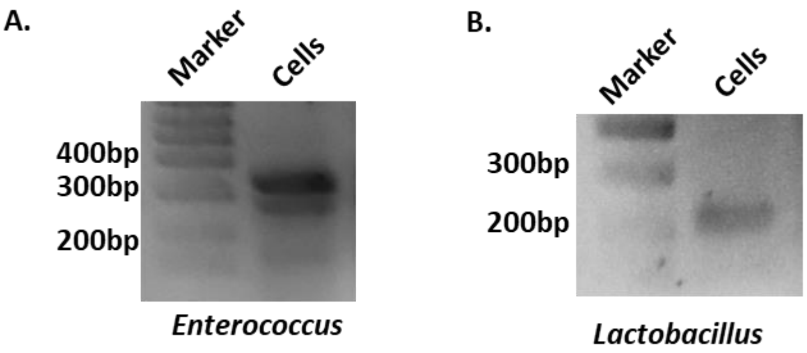


**Fig S4: Bacterial commensals are present within the cells of the Peyer's patches.** (A) *E. faecalis* identified in cells of Peyer's patches upon PCR amplification of isolated genomic DNA with *E. faecalis* specific primers. (B) Identification of *L. rhamnosus* in the cells of Peyer's patches cells in a similar manner.

#### Supplementary Fig. 5

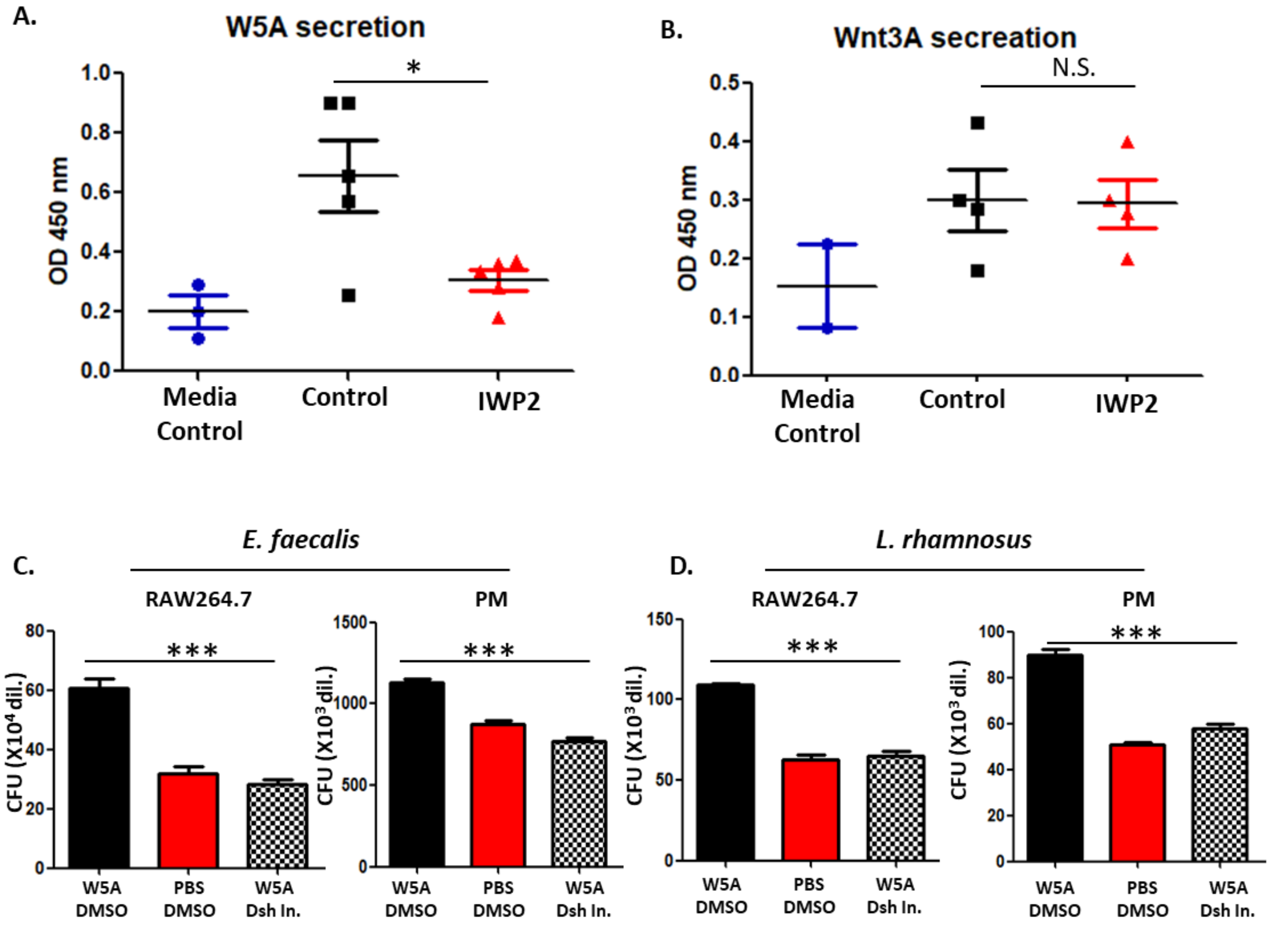


**Fig S5: Inhibition of Wnt5A signaling by Dsh inhibitor and IWP2.** (A) and (B) Application of 0.05µM IWP2 inhibited the secretion of Wnt5A (W5A) but not Wnt3A (W3A) from total cells of Peyer's patches as observed by indirect ELISA (n=5, n=4). (C) Application of 15µM of Dsh inhibitor in combination with rWnt5A during 6hr pre-incubation led to inhibition of internalization of *E. faecalis* by both RAW264.7 and peritoneal macrophages. The increase in internalization by Wnt5A signaling (Wnt5A-DMSO vs. PBS-DMSO) was nullified by Dsh (Dishevelled) inhibitor indicating that Dsh is a Wnt5A signaling intermediate (n=6 in RAW264.7, n=4 for PM). PBS and DMSO are vehocle controls for Wnt5A and the inhibitor respectively. (D) Application of Dsh inhibitor had a similar effect on *L. rhamnosus* internalization by RAW264.7 and peritoneal macrophages (n=4). "n" represents number of experiments. Data represented as mean ± SEM and $p \leq 0.05$ was considered as significant statistically. Significance was represented by * in the following manner: * $p \leq 0.05$, ** $p \leq 0.005$, *** $p \leq 0.0005$.

#### Supplementary Fig. 6

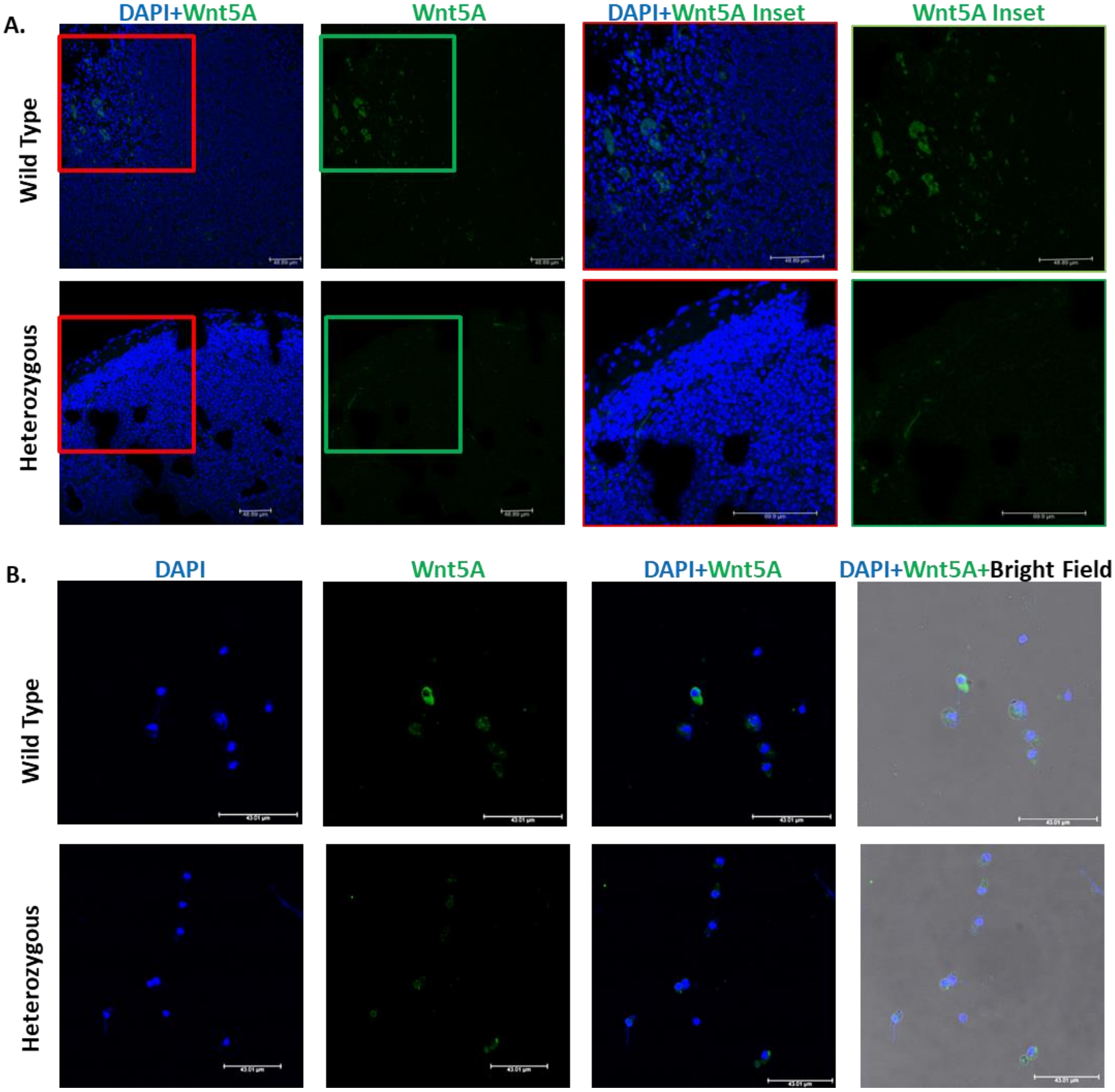


**Fig S6: Wnt5A expression is lower in Peyer's patches of Wnt5A heterozygous mice as compared to those of the wild type counterparts.** (A) Confocal microscopy of paraffin section of Peyer's patch tissue shows lower expression of Wnt5A (green) in heterozygous mice (bottom) as compared to wild type (n=2). (B) Wnt5A expression (green) in Peyer's patches of heterozygous mice were found to be lower than wild type counterpart even at the cellular level (n=2). "n" represents the number of mice in each group.

#### Supplementary Fig. 7

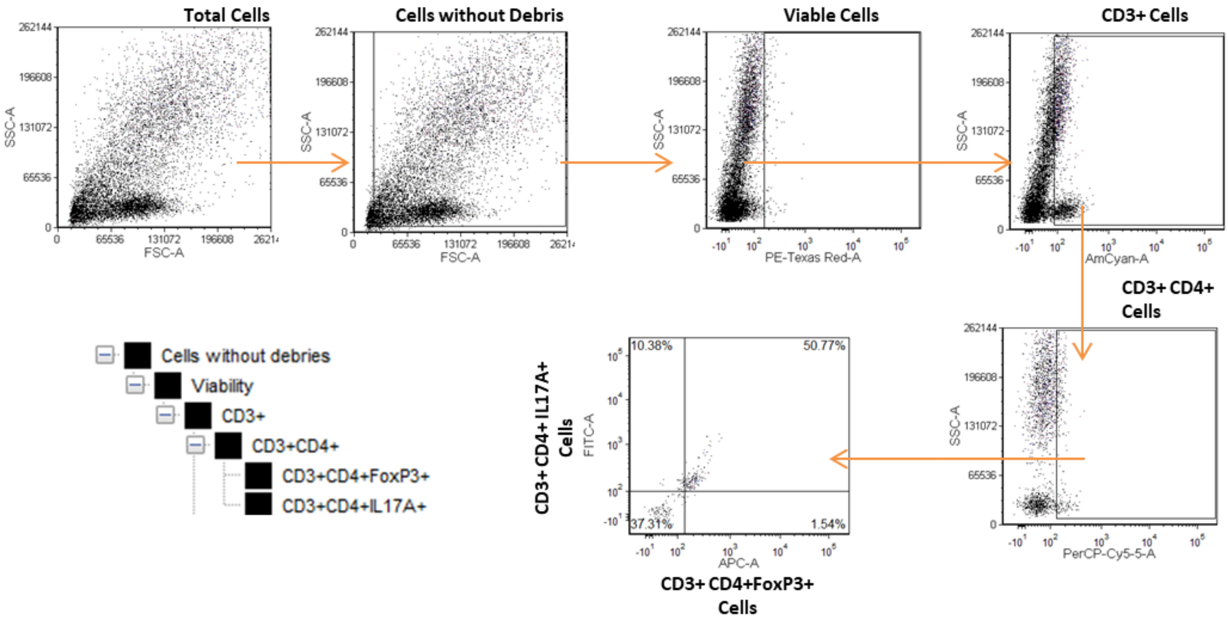


**Fig S7: Detection of IL17A+ and FoxP3+ T cell population.** Gating strategy for detection of IL17A+ and FoxP3+ T cell population shows reads with very low FSC and SSC (debris), which were ignored. Viable cells were selected out from total cell population (- debris). Within the viable population CD3+ cells were gated to identify the T cell population, following which CD4+ population was gated. IL17A+ and Fox P3+ population was then identified from CD3+CD4+ population via quadrant. Placement of the quadrant was done on the basis of unstained total population.

#### Supplementary Fig. 8

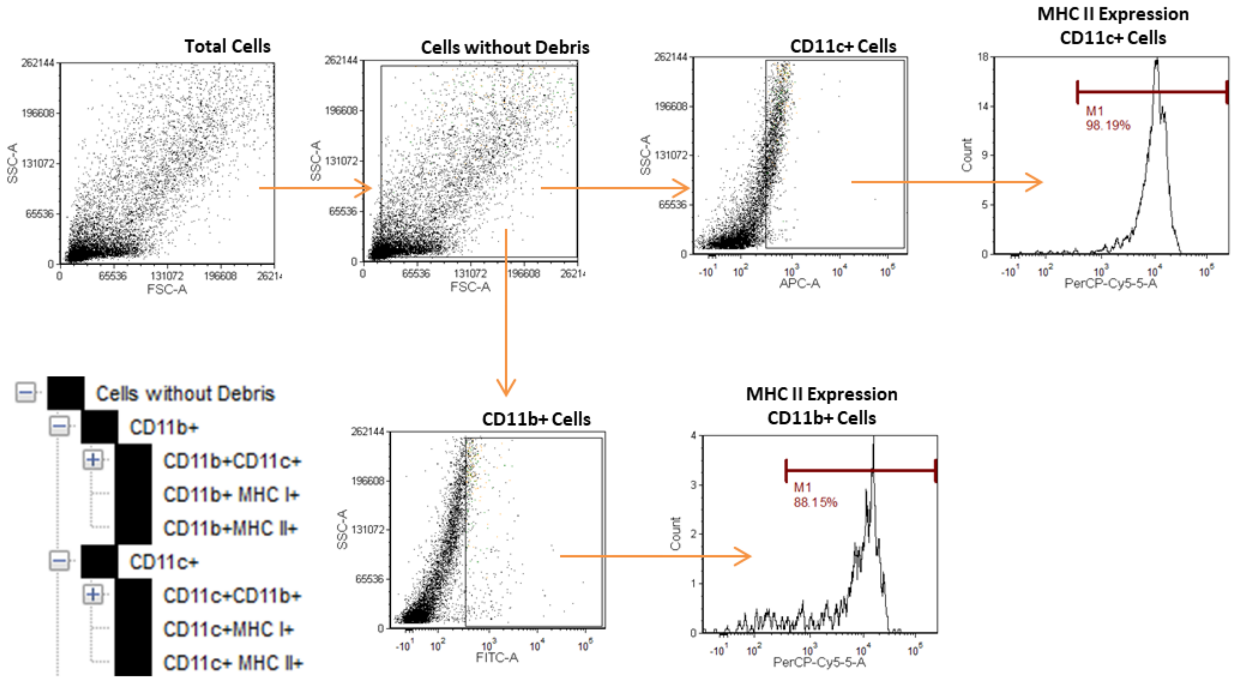


**Fig S8: Detection of MHC II expression in CD11b+ or CD11c+ phagocytes.** Gating strategy for observation of MHC II surface expression in CD11c+ and CD11b+ cells. Reads with very low FSC and SSC were ignored as noise, following which cells were gated for identification of CD11b+ or CD11c+ population. Histograms were then generated from each set of cells to observe intensity of MHC II expression within each population. Marker gates used for identification were based on unstained population.

#### Supplementary Fig. 9

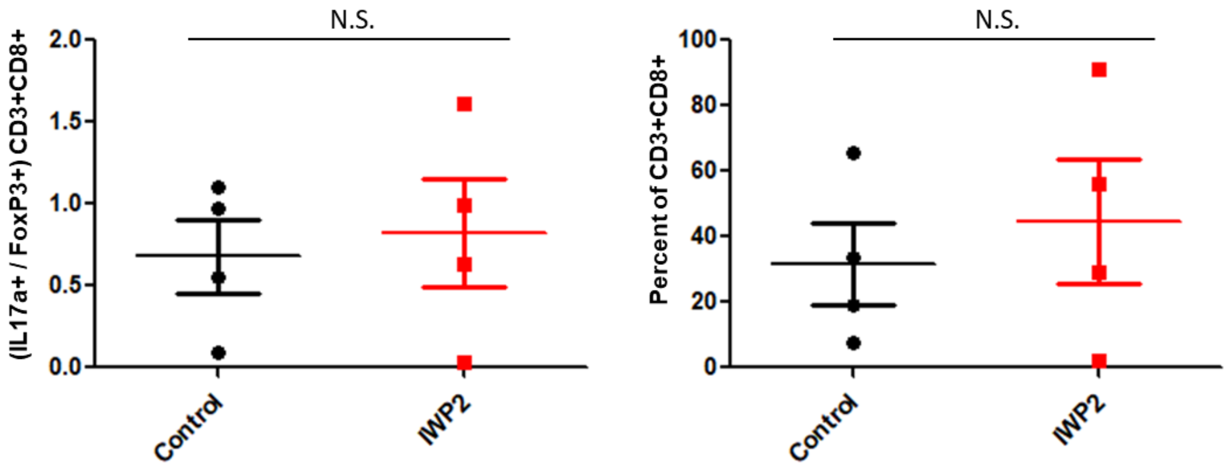


**Fig S9: Wnt5A mediated regulation of gut bacterial colonization did not have significant effect on CD8 T cell.** No significant effect of IWP2 treatment was observed on the IL17A+ FoxP3+ cells as and IL17A+FoxP3+ compartments of CD8+ T cells as represented in the graphs (n=4) "n" represents number of mice in each group. Data represented as mean ± SEM and p≤ 0.05 was considered as significant statistically. Significance was represented by * in the following manner: * p≤ 0.05, ** p≤ 0.005, *** p≤ 0.0005.
